# HIV-1 Reprograms CD4 T Cell Responses by Impairing Antigen-specific Communication with Dendritic Cells

**DOI:** 10.1101/2025.08.14.670333

**Authors:** Katharina Morath, Bettina Stolp, Jakob Rosenbauer, Karsten Mahnke, Michael Platten, Oliver T. Fackler

## Abstract

HIV-1 infection causes general dysfunction of adaptive immune cells that persists even under therapy but the underlying mechanisms remain elusive. Antigen-specific interactions of the main target cells of HIV, CD4 T cells, with dendritic cells (DCs) orchestrate global T cell responses and convey help to CD8 T cells. Here we report that HIV-1, by virtue of its pathogenesis factor Nef, impairs activation and transcriptionally reprograms CD4 T cells to dampen Th1 differentiation in response to antigen-specific stimulation by DCs. These alterations also disrupt functional communication to DCs to reduce DC activation and limit Th1 helper cytokine production. Mechanistically, Nef achieves this modulation of antigen-specific CD4 T cell function by reducing T cell surface levels of CD4. These results define modulation of CD4 T cell-DC communication as pathogenic principle by which HIV-1 disrupts adaptive immunity and emphasize the direct role of CD4 in immune cell communication.

**One Sentence Summary:** HIV-1 suppresses Th1 polarization by disrupting the communication between CD4 T cells and Dendritic cells through cell surface CD4 downregulation by the viral pathogenesis factor Nef.

## INTRODUCTION

Untreated infection with HIV-1 causes complex immunodeficiencies that prevent the adaptive immune system from protecting people living with HIV (PLWH) from opportunistic infections and thus the development of the acquired immunodeficiency syndrome (AIDS). HIV-1 replication can be efficiently controlled by livelong treatment with antiretroviral therapy (ART). However, emerging ART resistant virus variants require alternative treatment strategies and ART is often not available in areas of the global south most affected by the pandemic, a problem that has been potentiated by the recent rupture in ART supply (*1, 2*). Moreover, key aspects of HIV pathogenesis such as chronic inflammation and residual immune cell dysfunction persist even under ART and the mechanisms driving these immunopathologies are not well understood. AIDS thus remains a global health burden especially for developing countries (*3*). A central problem in AIDS pathogenesis is that adaptive immune responses are impaired in PLWH. This reflects on one hand immune evasion mechanisms enabled by the genetic plasticity of HIV that drives the selection of viral proteins with low immunogenicity at fully preserved functionality. In addition, HIV exerts active mechanisms that contribute to global immunodeficiency and result in dysfunction of B cells and CD8 T cells in PLWH. In particular, CD8 T cell dysfunction is a hallmark of HIV infection: while initial CD8 T cell responses efficiently reduce circulating virus, they are driven into exhaustion and senescence over time. These dysfunctions can be reverted to some degree under ART but are not completely restored even after years of treatment (*4–7*).

B cell and cytotoxic T cell responses require CD4 T cell help and the depletion of CD4 T cells likely accounts for broad immune dysfunction at late stages of disease. This mechanism however does not explain the impairment of immune function during the asymptomatic phase or under ART where CD4 T cell counts are normal, suggesting that HIV interferes with these responses by directly impairing CD4 T cell function (*4, 7, 8*). The existence of such mechanisms is indicated by the altered CD4 T cell differentiation patterns (e.g. shift from Th1 to effectorless Th0 CD4 T cells) and cytokine signatures (e.g. hyperproduction of the immune regulatory cytokine IL-7) observed in PLWH (*9–13*) as well as general chronic inflammation, but the mechanisms driving these dysfunctions remain elusive.

To exert helper functions, CD4 T cells engage in close and antigen-specific physical contacts with antigen-presenting cells referred to as immune synapse (IS). IS formation drives CD4 T cell activation and differentiation but also triggers the production of key helper cytokines (*14–18*). While B cell function is regulated by direct interaction with cognate CD4 T cells, CD8 T cell responses are regulated by CD4 T cell interaction with Dendritic Cells (DC) that orchestrate global T cell responses and mediate CD4 T cell help to CD8 T cell responses (*19–27*). Importantly, ISes of CD4 T cells with B cells or DCs are fundamentally different regarding their stability, architecture and potency of T cell activation (*28–33*) and mechanisms that govern these distinct interaction modes and T cell stimulation efficiencies are poorly understood. Building on our previous finding that the HIV-1 accessory protein and pathogenicity factor Nef impairs antibody response by abrogating functional CD4 T cell interactions with and help to B cells *in vivo* (*34*), we assessed here whether HIV alters the ability of CD4 T cells to communicate with DCs towards the mounting of CD8 T cell responses. We identify that HIV, by virtue of its pathogenesis factor Nef, uses downregulation of cell surface CD4 as a unique mechanism to disrupt antigen-specific CD4 T cell activation, differentiation and communication with DCs.

## RESULTS

### HIV-1 Nef interferes with early antigen-specific CD4 T cell-DC interaction *ex vivo*

Antigen-specific interactions of CD4 T cells with DCs not only drive potent T cell activation and differentiation into T-bet+ Th1 cells but also provide important help to the DCs, driving their activation and production of Th1 polarizing cytokines such as IL-12 that ultimately support mounting of CD8 T cell responses (Fig. 1A). Depending on the experimental set-up, the HIV pathogenesis factor Nef has been reported to induce or impair experimental CD4 T cell activation (*35–42*). These studies analyzed CD4 T cell responses to chemical or antibody-mediated stimulation or used superantigen presentation, which does not assess bi-directional signaling between antigen presenting and T cell. To investigate the impact of HIV-1 Nef on antigen-specific, bi-directional communication between CD4 T cells and DCs, we activated murine OT-II.2 CD4 T cells that recognize the OVA323-339 peptide in the context of MHC class II I-Ab expression and used MLV-based transduction to induce expression of HIV-1 SF2 Nef to levels that are equivalent to that of HIV-1 infection of primary human CD4 T cells (*43*). Transduced CD4 T cells were sorted based on their expression of a truncated version of the human NGFR and co-cultured with bone-marrow derived DCs that were differentiated with GM-CSF and either left in an immature state (iDC) or matured overnight by addition of LPS (mDC). iDCs and mDCs were pulsed with increasing concentrations of OVA323-339 immediately before co-culture at approximately 1:4 DC:T cell ratio (Fig. 1B) and cytokine production, activation and proliferation of CD4 T cells was assessed at various timepoints. Expectedly, production of IL-2 and IFNγ by CD4 T cells was induced in an antigen dose-dependent manner and was more efficient when antigen was presented by mDCs than by iDCs (Fig. 1C and D and Fig. S1A-C). Importantly, this cytokine induction was significantly impaired in Nef-expressing CD4 T cells (Nef) compared to CD4 T cells transduced with an empty control vector (Ctrl) (4.0±1.3% IL-2+IFNγ+ cells for Nef versus 13.9±3.0%, for Ctrl with iDC and 8.4±1,3% versus 25.1±0.6% for mDC co-culture with 1000 nM OVA), resulting in the lack of a response at lower and a reduced response at the highest dose of antigen used. This Nef-mediated impairment of antigen-specific activation was also reflected by a reduced induction of the T cell activation marker CD69 at low antigen doses, an inhibition that was overcome at the highest antigen concentration used (CD69 GeoMean of 4594.1±1037.9 for Nef versus 6767.0±1683.5 for Ctrl iDC co-cultures and 7432.6±1276.7 versus 9210.6±1478.3 for mDC co-cultures at 1000 nM OVA, respectively) (Fig. 1E and F). In contrast, activation-induced proliferation of CD4 T cells was not significantly impaired by Nef (Fig. 1G and H). Nef thus impairs CD4 T cell activation upon engagement by antigen-presenting DCs but these effects primarily act on early steps of T cell activation and can be overcome by saturating antigen doses over time.

**Fig. 1.**
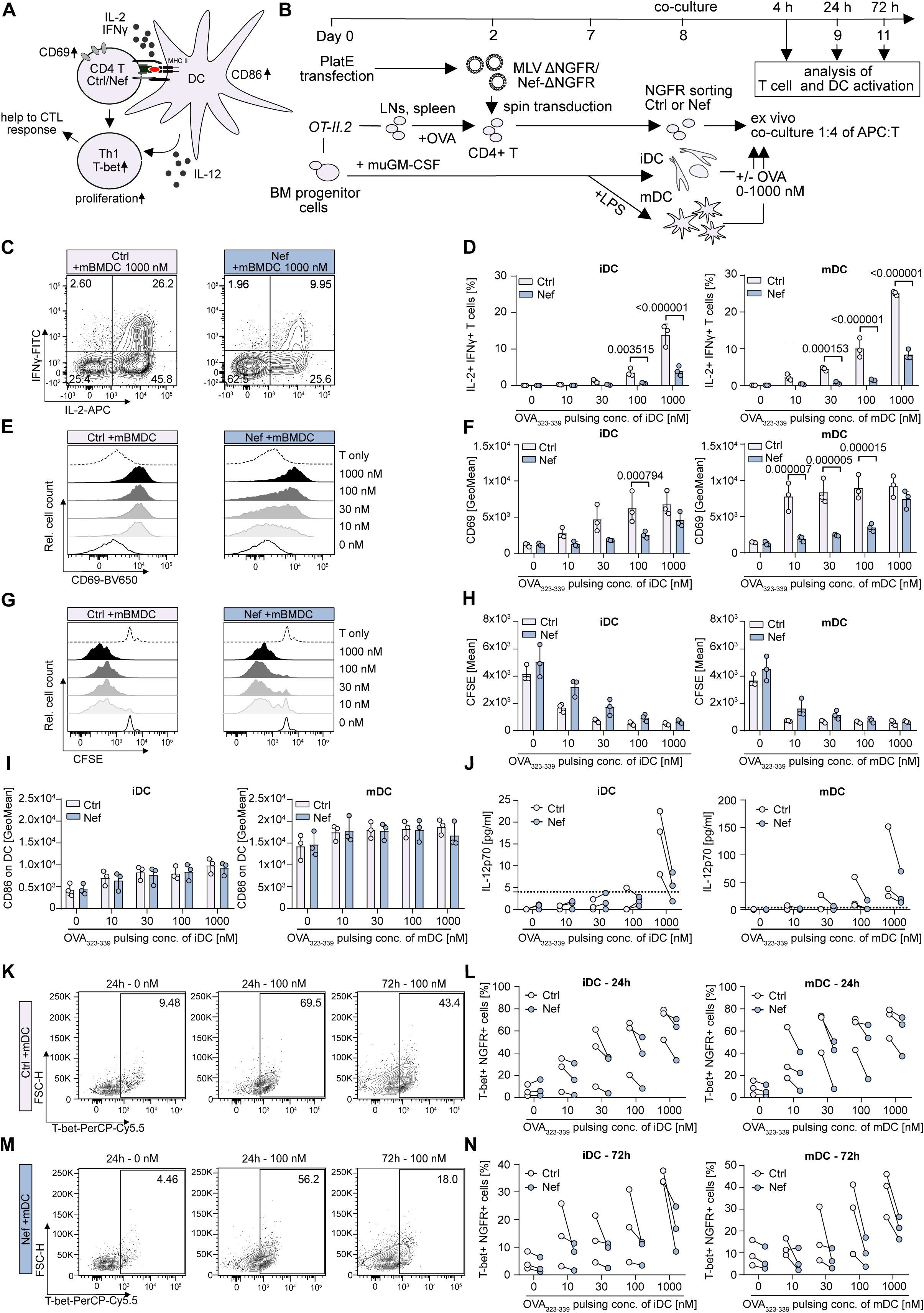
HIV-1 Nef impairs early cognate CD4 T cell-DC interactions and Th1 differentiation *ex vivo*. **(A)** Schematic of CD4 T cell-DC interaction. **(B)** Experimental set-up. **(C)** Representative flow cytometry plots of Ctrl or Nef cells showing IL-2 and IFNγ production at 4.5h upon cognate interaction with mDC. **(D)** Quantification of C for indicated conditions. **(E)** and **(G)** Representative histograms showing CD69 expression at 24h (E) or CFSE dilution at 72h (G) for indicated conditions. **(F)** and **(H)** Quantification of E and G. **(I)** Quantification of CD86 expression on DCs at 24h of co-culture. **(J)** IL-12p70 concentrations in supernatants of co-cultures at 24h measured by ELISA. Dashed line indicates the detection limit. **(K)** and **(M)** Representative flow cytometry plots showing intranuclear T-bet signal in Ctrl or Nef cells in DC co-culture at 24 or 72h with or without antigen as indicated. **(L)** and **(N)** Quantification of K and M. D, F, H and I. Shown are means with SD from three independent experiments, each symbol represents the mean of biological triplicates per experiment. J, L and N. Shown is data from three independent experiments, with matched data per experiment indicated by connecting lines between Ctrl- and Nef cell co-cultures. Each symbol represents the mean of technical duplicates (J) or biological triplicates (L and N). Statistical significance in D, F, I, L and N was calculated using multiple t-test, Wilcoxon test was used in H and J. p-values are indicated for significant differences between Ctrl and Nef conditions.

We next investigated how expression of HIV-1 Nef in CD4 T cells affects activation and maturation of interacting DCs. Cognate interactions with CD4 T cells induced maturation of iDCs, as assessed by an increase in CD86 surface expression at higher antigen concentration, but this effect was unaffected by Nef (Fig. 1I and Fig. S1D and E). In contrast, production of IL-12p70 by DCs, which was best appreciable at high antigen doses, was reduced upon interaction with Nef as compared to Ctrl CD4 cells (Fig. 1J). Nef thus impairs the ability of CD4 T cells to stimulate DC cytokine production. Since both, IL-12 and IFNγ, are essential for directing CD4 T cell differentiation towards Th1 responses, we next assessed intracellular levels of the Th1 marker T-bet in CD4 T cells at 24 and 72h of co-culture with DCs. While T-bet was efficiently induced in Ctrl CD4 T cells in a dose-dependent manner by iDCs and mDCs (up to 79.2% and 46.0% T-bet positive cells at 24h or 72h) (Fig. 1K-N and Fig. S1F-H), T-bet expression was reduced in Nef CD4 T cells at both timepoints, even at the highest antigen concentration. We conclude that in these *ex vivo* co-cultures, Nef expression in CD4 T cells impairs their activation by cognate DCs and limits mutual cytokine production and that these effects are associated with the suppression of Th1 polarization of Nef CD4 T cells. These results suggest that the direct effects of Nef expression in CD4 T cells affect DC function which in turn feeds back to CD4 T cells and collectively block Th1 polarization.

### HIV-1 Nef prolongs cognate CD4 T cell interactions with DCs within lymph nodes *in vivo*

Physiologically, CD4 T cell interactions with DCs take place in secondary lymphoid organs and spatial architecture as well as the unique cellular environment of these tissues provide important cues for T cells and DCs. To account for these environmental factors, we next assessed CD4 T cell activation and proliferation *in vivo*. To this end, OT-II.2 CD4 T cells (CD45.2) were sorted and stained with Celltrace violet (CTV) prior to i.v. injection into recipient SMARTA animals (CD45.1). One day later, matured DCs were loaded with different OVA concentrations (0-100 nM) and injected s.c. above the ankle of front or hind legs and allowed to migrate to the draining lymph node to create distinct reaction centers with increasing antigen availability within each recipient animal. One day later, FTY720 was administered to block T cell egress and 24 or 72h later, draining lymph nodes of the DC injection sides (axillary and brachial lymph nodes for front leg, popliteal lymph nodes for hind leg injections) were isolated and processed for flow cytometry analysis (Fig. 2A). As expected, only a small population of CD45.2 OT-II.2 CD4 T cells could be detected at 24 and 72h in draining lymph nodes in the presence of unpulsed DCs. This population markedly expanded until 72h for lymph nodes draining antigen-pulsed DCs (Fig. 2B and Fig. S2A) and CD69 expression of lymph nodes draining antigen-pulsed DCs increased in a dose-dependent manner already at 24h (Fig. 2C and E), followed by an increase in T cell proliferation at 72h (Fig. 2D and F). Likely reflecting the homing defect of Nef-expressing CD4 T cells (*43*), less Nef CD4 T cells than Ctrl CD4 cells were detected at 24 and 72h within lymph nodes (Fig. 2B and G and Fig. S2B). Importantly and in line with the results from the *ex vivo* co-cultures, Nef expression significantly reduced activation at 24h (rel. CD69 Median of 45.1% for Nef versus 93.4% for Ctrl and 66.3% versus 101.9% for 30 and 100 nM, respectively) of successfully homed CD4 T cells, but had no effect on general T cell proliferation. (Fig. 2E, F and G).

**Fig. 2.**
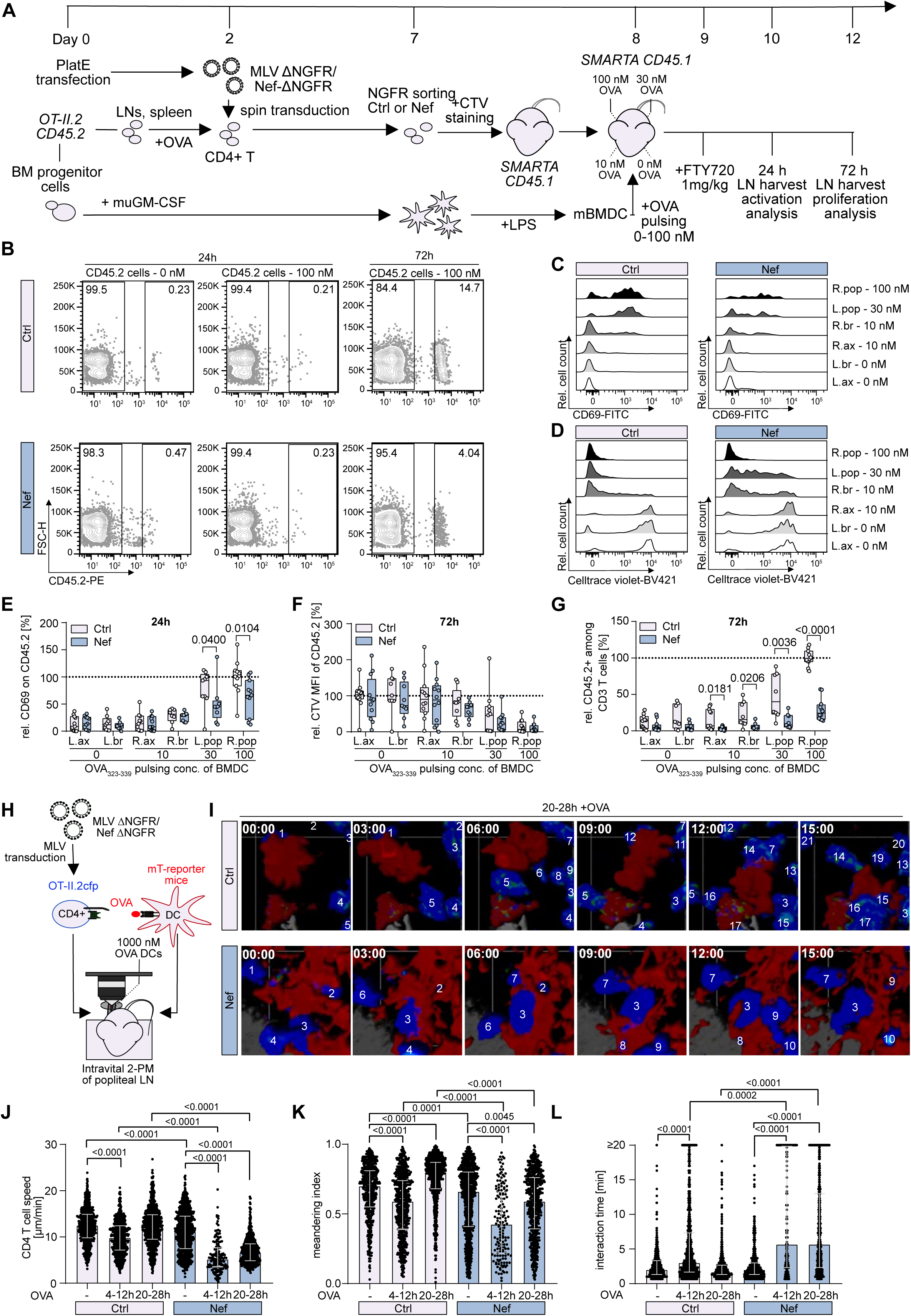
HIV-1 Nef impairs antigen-specific CD4 T cell-DC interaction *in vivo* and prolongs interaction times within draining lymph nodes. **(A)** Experimental set-up. **(B)** Representative flow cytometry plots of draining lymph nodes of CD45.1 SMARTA mice at 24h or 72h showing frequency of CD45.2 cells among total CD3+ cells for Ctrl or Nef conditions. **(C)** and **(D)** Representative histograms of CD69 signal at 24h and CTV signal at 72h for indicated conditions. **(E)** and **(F)** Quantification of C and D. **(G)** Quantification of CD45.2 cells among CD3+ cells at 72h. E, F and G. Shown are box and whisker plots with median, min and max range for n=4 individual experiments with 2-3 animals per group per experiment. Each dot represents data from one animal per condition. **(H)** Experimental set-up for intravital 2PM analysis at early (4-12h) and late (20-28h) timepoints. **(I)** Representative still images with timestamps from 2PM movies. Numbers mark individual T cells that entered the field of view over time. **(J)**, **(K)** and **(L)** Quantification of mean migration speed (J) and meandering index (K) of individual T cell tracks and interaction times of individual T cells with DCs (L) as shown in I. Shown are medians with interquartile range for data obtained from 1-2 independent experiments per condition with a total of 3-6 animals and 6-13 movies per condition. Each dot represents one track. E, F and G. Statistical analysis was performed using Mann-Whitney test in E, F and G and Kruskal-Wallis test in J, K and L. p-values are indicated where statistically significant.

Since our results suggested that Nef interferes in particular with early steps of T cell - DC interactions, we next employed intravital 2-Photon microscopy to visualize the dynamics of T cell - DC interactions within secondary lymphoid organs. Transduced and sorted OT-II.2cfp CD4 T cells and mDCs from mTomato-reporter mice were loaded with 1000 nM OVA and injected intravenously, i.e. subcutaneously into the hind hock of recipient animals (Fig. 2H). CD4 T-DC interaction dynamics are known to alter over time. To assess these various interaction modes, we performed imaging of cell interactions within the popliteal lymph node at 4-12h and 20-28h (i.e. 20-28 and 36-44h post DC injection). These analyses revealed frequent and dynamic interactions of T cells (blue) with DCs (red) within lymph nodes and changes in interaction behavior in dependence of antigen and imaging timepoint (Fig. 2I, Fig. S2C and movies S1-S6). As expected, the migration speed of Ctrl CD4 T cells decreased significantly from a median of 12.5 to 9.8 µm/min in the presence of antigen-presenting DCs at 4-12h, paralleled by a decrease in the directionality of movement (meandering index 0.70 without OVA versus 0.59 with OVA at 4-12h) due to antigen-specific interaction with DCs (Fig. 2J-L). Normal motion was resumed at 20-28h and conversely, directional movement increased, which likely reflects the short engagement with antigen-bearing DCs (Fig. 2J-L). Similar to Ctrl cells, Nef CD4 T cells arrested migration upon encounter with antigen-pulsed DCs. However, T cell-DC interactions were significantly longer compared to Ctrl CD4 T cells at both early and late timepoints (median of 5.67 min Nef CD4 T cells vs. 3 and 1.67 min for Ctrl cells) (Fig. 2J-L). Collectively, these results suggest that Nef manipulates and prolongs interaction dynamics between CD4 T cells and DCs *in vivo* which is associated with a reduced antigen-specific activation response of CD4 T cells and suboptimal licensing of DCs.

### HIV-1 Nef expression in T cells dampens pro-inflammatory cytokine production upon cognate DC interaction

The above results revealed that modulation of CD4 T cell-DC interactions by HIV-1 Nef *in vivo* and *ex vivo* is associated with less efficient Th1 differentiation. To gain mechanistic insight, we characterized the breadth and magnitude of functional changes and assessed the secretion of a broad range of cytokines and chemokines in the supernatant of *ex vivo* CD4 T cell – DC cultures (Fig. 3A). Basal cytokine production of Ctrl CD4 T cells in the absence of antigen was generally low but pulsing of iDCs or mDCs with antigen prior to co-culture triggered the production of a broad range of cytokines (Fig. 3B). Among the analysed factors, three major groups could be distinguished; (i) factors associated with antigen-specific interaction of T cell and DCs, (ii) factors associated with DC maturation as well as antigen-specific interaction and (iii) factors not induced by antigen-specific interaction or only detectable at very low levels (Fig. 3B, Fig. S3, Table S1 and Data S1). Low basal, antigen-independent cytokine expression was only moderately altered in the presence of Nef CD4 T cells and these effects varied between experiments (Fig. 3B). In contrast, production of many cytokines in the presence of antigen was reproducibly impaired by Nef expression in CD4 T cells (Fig. 3B, C). While factors unrelated to antigen-specific interaction, such as CCL2, M-CSF and CXCL10, were unaffected, Nef consistently suppressed cytokines induced by antigen-specific interaction of T cells and DCs irrespective of the DC maturation status including IL-2, TNFα, IFNγ, IL-10, IL-3, IL-5, CCL3 and CCL4 (Fig. 3C-K). These results confirm that Nef limits CD4 T cell activation upon cognate DC interactions and suppresses the expression of activation-induced T helper cytokines such as IL-2, TNFα and IFNγ to generate an anti-inflammatory microenvironment that is unfavorable for Th1 polarization (Fig. 3D-F). In addition, cytokines deregulated by Nef such as IL-10, IL-3 and IL-5 exert important roles in the activation of DCs and cytotoxic CD8 T cells (Fig. 3G-I) (*44–49*). Further, CCL3 and CCL4 promote the ability of APCs to support Th1 polarization, recruit naïve CD8 T cells to licensed DC/CD4 T cell clusters *in vivo* (*50, 51*), and are strongly linked to the anti-HIV activity of CD8 T cells (*52, 53*) (Fig. 3J and K). These Nef-mediated deregulations were mild when considering individual cytokines but resulted in a broad change of the complex microenvironment that collectively is suboptimal for Th1 and CD8 T effector cell generation.

**Fig. 3.**
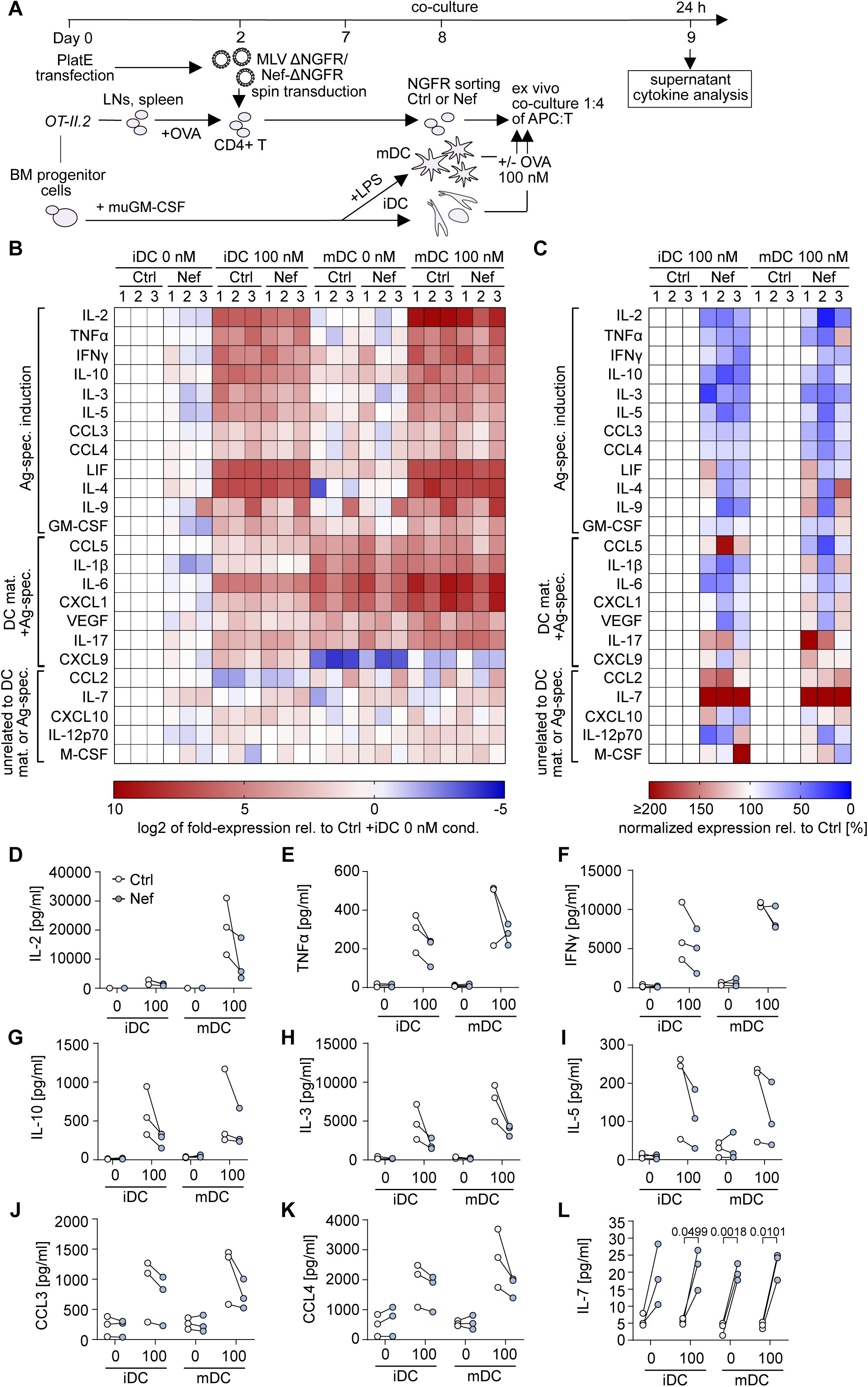
HIV-1 Nef alters the local microenvironment upon antigen-specific interaction of CD4 T cells with DCs. **(A)** Experimental set-up. **(B)** Heatmap summarizing cytokine levels in supernatants of co-cultures from three independent experiments (1, 2, 3) showing log2-fold changes in cytokine concentrations relative to the Ctrl +iDC 0 nM condition per cytokine. **(C)** Heatmap showing relative cytokine production in Nef co-cultures with antigen in % relative to Ctrl cells. **(D)**-**(L)** Absolute cytokine concentrations for IL-2, TNFα, IFNγ, IL-10, IL-3, IL-5, CCL3, CCL4 and IL-7. Values are presented in pg/ml. Shown are values from three independent experiments, each symbol represents data from one experiment and lines connect matched data for Ctrl or Nef T cell co-cultures per experiment. Statistical analysis was performed using Wilcoxon test for IL-2 and TNFα and paired t-test for IFNγ, IL-10, IL-3, IL-5, CCL3, CCL4 and IL-7. p-values are indicated where statistically significant differences between Ctrl and Nef conditions were observed.

IL-7 represented a notable exception of a cytokine that was strongly induced rather than suppressed upon Nef expression and deregulation occurred in the absence as well as presence of antigen (Fig. 3C and L). IL-7 is a homeostatic cytokine that has been associated with viral persistence and facilitation of HIV-1 infection in resting T cells (*54–56*) and its induction by Nef may thus contribute to the boost of virus replication provided by the pathogenesis factor *in vivo* (*57, 58*). Nef thus alters the cytokine milieu to generate a microenvironment that suppresses cellular immunity while favoring virus spread.

### Nef expression in CD4 T cells reprograms the transcriptional response of CD4 T cells and DCs to cognate interactions

To unravel the breadth of functional changes imprinted by Nef on CD4 T cell – DC communication, we next performed scRNAseq analysis on 4 and 24h co-cultures of OT-II.2 CD4 T cells and DCs (Fig. 4 and Fig. S4-S7). Cell type-specific gene expression profiles allowed to identify T cell and DC populations (Fig. 4A) and subgroups that were submitted to more detailed analysis and clustering (Fig. 4B and C, Fig. S4A and B). Expectedly, we observed global gene expression changes between the 4h and 24h timepoints post cognate interaction in both T cells and DCs (Fig. S4C-F). These included TCR and cytokine signaling genes in T cells and were most pronounced in the activated ‘cytokine and MYC T’, ‘proliferating T_1’ and ‘MYC and translation T’ cell subsets (Fig. S5A-E and V). In DCs, motility and maturation-associated genes were broadly induced (Fig. S5N-U and W). Although the general cluster distribution was similar between Ctrl and Nef CD4 T cell co-cultures, Nef-induced alterations in gene expression at both early and late timepoints resulted in the expansion or reduction of specific subsets of T cells (Fig. 4B) and DCs (Fig. 4C). Pathway analysis revealed that in CD4 T cells, the most pronounced effects of Nef early post stimulation included the repression of cytokine signaling-, Rho GTPase-, receptor tyrosine-kinase- and NFkB-related pathways, which was reflected by reduced *Ifng*, *Il2* and *Stat5b* expression and paralleled by reduced expression of the Th1-associated genes *Tbx21* and *Runx3* suggesting activation and Th1 polarization is suppressed on a transcriptional level by Nef. In turn, pathways involved in metabolic and translation control were enriched for Nef cells at this timepoint, indicating higher protein biosynthesis (Fig. 4D, Fig. S6A-E). Consistent with an impairment of CD4 T cell activation, translation associated genes (such as *Rpl32* and *Rpl10*) that dominated the antigen-specific response at 24h in Ctrl T cells were strongly reduced in Nef cells at that timepoint (Fig. 4E, Fig. S5F and G and Fig. S6F, G and N). Conversely, Nef expressing cells maintained high expression of a small group of genes that were downmodulated in an antigen-specific manner in Ctrl cells at 24h. Interestingly, these included the genes encoding the Nef protein interaction partners Elmo1 (*Elmo1*) and Dock2 (*Dock2*) that, together with Nef, activate the Rho GTPase Rac1 to induce TCR-independent T cell activation (Fig. S5H-M and Fig. S6H-M) (*59*). Consequently, maintained expression was associated with enrichment of signal transduction and second messenger pathways at 24h, indicating a delayed activation response (Fig. 4E, Fig. S6N). To investigate the possibility that Nef cells preferentially differentiated into other CD4 T cell subtypes than Th1, such as Th17, Th2 or Treg cells, we analysed expression of transcription factors RORγt (*Rorc*), GATA3 (*Gata3*) and Foxp3 (*Foxp3*) but observed no enhanced alternative induction of these transcription factors (see deposited scRNAseq dataset). Collectively, these results show that Nef impairs the transcriptional response of CD4 T cells upon cognate interaction with DCs and promotes TCR-independent activation pathways.

**Fig. 4.**
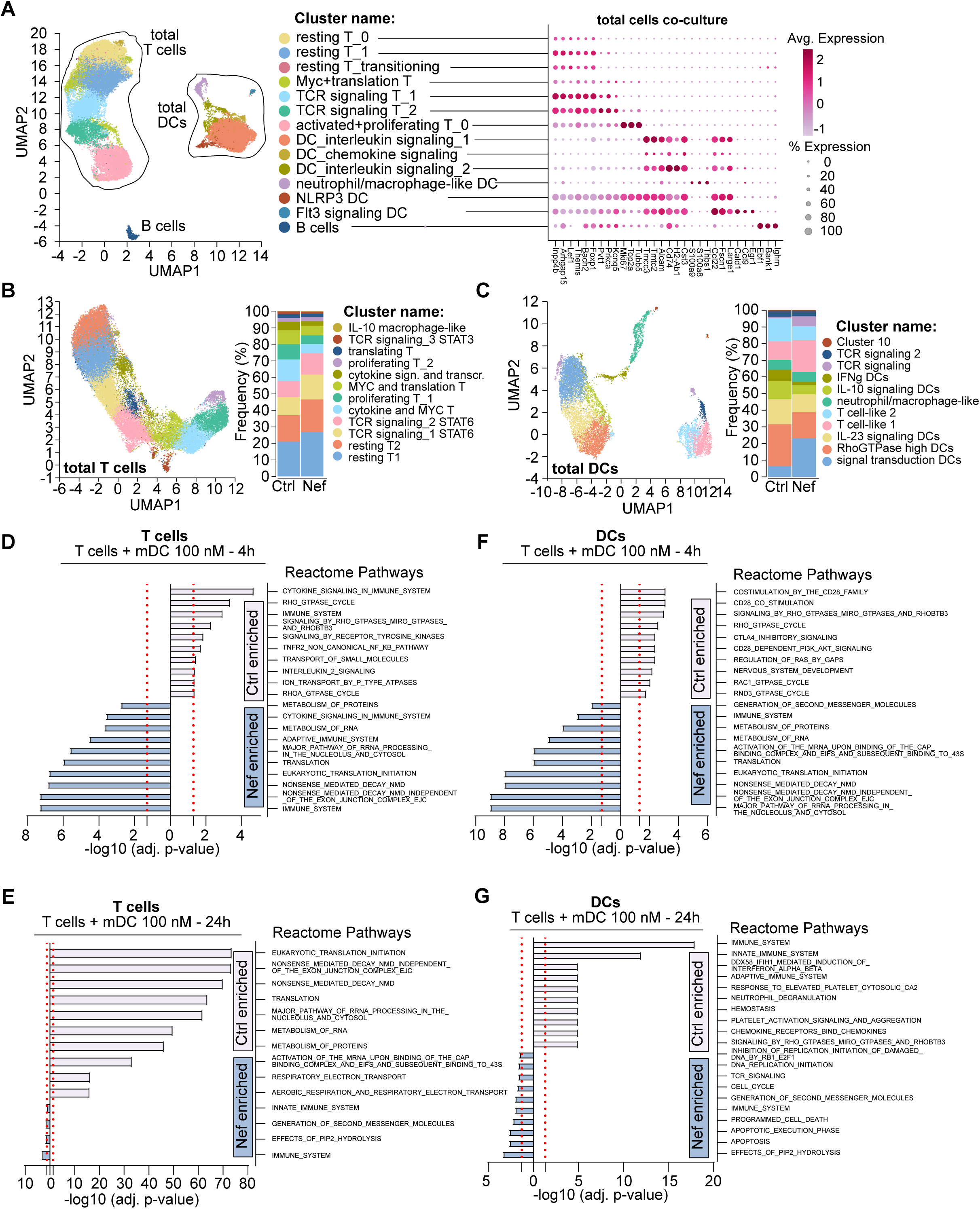
HIV-1 Nef manipulates transcriptional responses in both CD4 T cells and DCs upon antigen-specific interaction. **(A)** UMAP of total cells of CD4 T cell co-cultures with DCs (left) resulting from scRNAseq analysis and dot plot showing differential marker gene expression of the identified clusters (right). Cluster annotation was performed with the Enrichr tool using the top 50 differentially expressed genes per cluster. **(B)** and **(C)** UMAPs of T cell (B) or DC (C) populations subgated from total cells based on gene expression profiles with subcluster distribution in stacked bargraphs for Ctrl or Nef conditions, respectively. **(D)**-**(G)** Pathway enrichment analysis based on top 100 differentially expressed genes between Ctrl or Nef T cell co-cultures with antigen-pulsed mDC at 4h (D and F) or 24h (E and G) based on the Mouse Reactome 2024 database in MSigDb. Shown is data for either T cells (D and E) or DCs (F and G). Shown are –log10 values of adjusted p-values. Red dashed lines represent adjusted p-value cutoff of 0.05.

Reflecting these alterations in CD4 T cell function by the viral pathogenesis factor, DCs displayed impaired antigen-dependent upregulation of motility-co-stimulation- and maturation-associated genes at 4h post interaction with Nef CD4 T cells. This deregulation included *Alcam,* a regulator of IS formation with T cells, *Fscn1, Cd80, Cd86, CD40* and *Arhgap26* that are involved in DC co-stimulation, maturation and motility, and *Rictor* and *Yes1* that participate in signaling and gatekeeping of DC activation (Fig. S5N-U and Fig. S7A-H) (*60, 61*). Consequently, pathway enrichment analysis revealed insufficient induction of CD28 co-stimulation and Rho GTPase signaling pathways at 4h compared to DCs co-cultured with Ctrl CD4 T cell (Fig. 4F). At later timepoints, DCs in antigen-specific co-culture with Ctrl CD4 T cells upregulated protein biosynthesis and immune system pathways while DCs co-cultured with Nef CD4 T cells showed delayed induction of GTPase and signaling pathways (Fig. 4G and Fig. S7I). Together, these results reveal that Nef expression in CD4 T cells alters the output of cognate communication with DCs that delays and tunes the responses of both cell types upon antigen-specific interaction by transcriptional reprogramming. In CD4 T cells, Nef tunes the transcriptional response towards a signature repressive for activation and Th1 polarization associated genes while transcription of genes associated with TCR-independent activation is enhanced. Conversely, DCs interacting with Nef CD4 T cells display suboptimal induction of maturation- and motility associated genes which is in line with the observed reduction of release of Th1 and CD8 T cell help-associated factors.

### Molecular determinants of Nef-mediated modulation of cognate CD4 T cell-DC communication

HIV-1 Nef is a multifunctional protein that optimizes viral replication and persistence by hijacking cellular pathways by virtue of its ability to recruit host cell proteins to multiple independent molecular surfaces (Fig. 5A). To define effector functions that are required for Nef disruption of T cell-DC interactions, we employed a panel of well-characterized HIV-1 SF2 Nef mutants in which individual protein interaction motifs are disrupted to allow segregation of different Nef effector functions (*62*). Disrupted interaction motifs included a di-leucine motif required to target the cellular endocytosis machinery for internalization of cell surface receptors (Nef LLAA mutant, (*63*)), a PxxP SH3 domain binding motif required for interference with cellular signaling and transport pathways (Nef AxxA, (*64*)), the interaction motif for the PAK2 kinase that allows Nef to interfere with host cell actin dynamics (Nef F195A, (*65, 66*)) and the SERINC5 antagonism motif (S5AM) required for antagonism of the restriction factor SERINC5 and disruption of CD4 T cell help to B cells (Nef A32-39, (*34, 67*)) (Fig. 5A and B). Analyzing these mutants for their ability to interfere with the T cell as well as the DC side of this immune cell communication revealed that the Nef F195A variant was as active as wild type Nef in reducing antigen-induced CD4 T cell cytokine secretion and activation (Fig. 5C, D, F, G and Fig. S8A-C, D, F-H, J, L-N) as well as IL-12 production by DCs (Fig. 5H and Fig. S8I and O). Interference with cellular actin dynamics is therefore dispensable for the modulation of CD4 T cell-DC communication by Nef. In contrast, the Nef LLAA and Nef A32-39 mutants displayed reduced inhibition of early cytokine production compared to Nef WT (mean of 22.30±1.05% for Nef LLAA and 17.9±1.0% for Nef A32-39 versus 9.0±0.7% IL-2+ cells for Nef WT cells and 38.4±0.3% for Ctrl cells for 100 nM co-cultures, respectively) (Fig. 5C and D and Fig. S8D and J), failed to inhibit CD4 T cell activation at 24h (CD69 Mean of 3950.3±419.2 and 2755.0±134.4 versus 3640.7±57.0 for Ctrl cells in 30 nM co-cultures), and their ability to reduce IL-12p70 production by DCs was reduced (Fig. 5F-H and Fig. S8A-C, F-I and L-O). Nef AxxA was partially active with intermediate effects on early and late T cell activation as well as DC cytokine production. Notably, the strongest loss of Nef interference with CD4 T cell – DC communication by the LLAA and A32-39 mutants was associated with a complete loss of the ability to downregulate cell surface levels of the HIV-1 entry receptor CD4 (Fig. 5E and Fig. S8E and K). The Nef-mediated impairment thus depends on two molecular surfaces that contribute to CD4 downregulation.

**Fig. 5.**
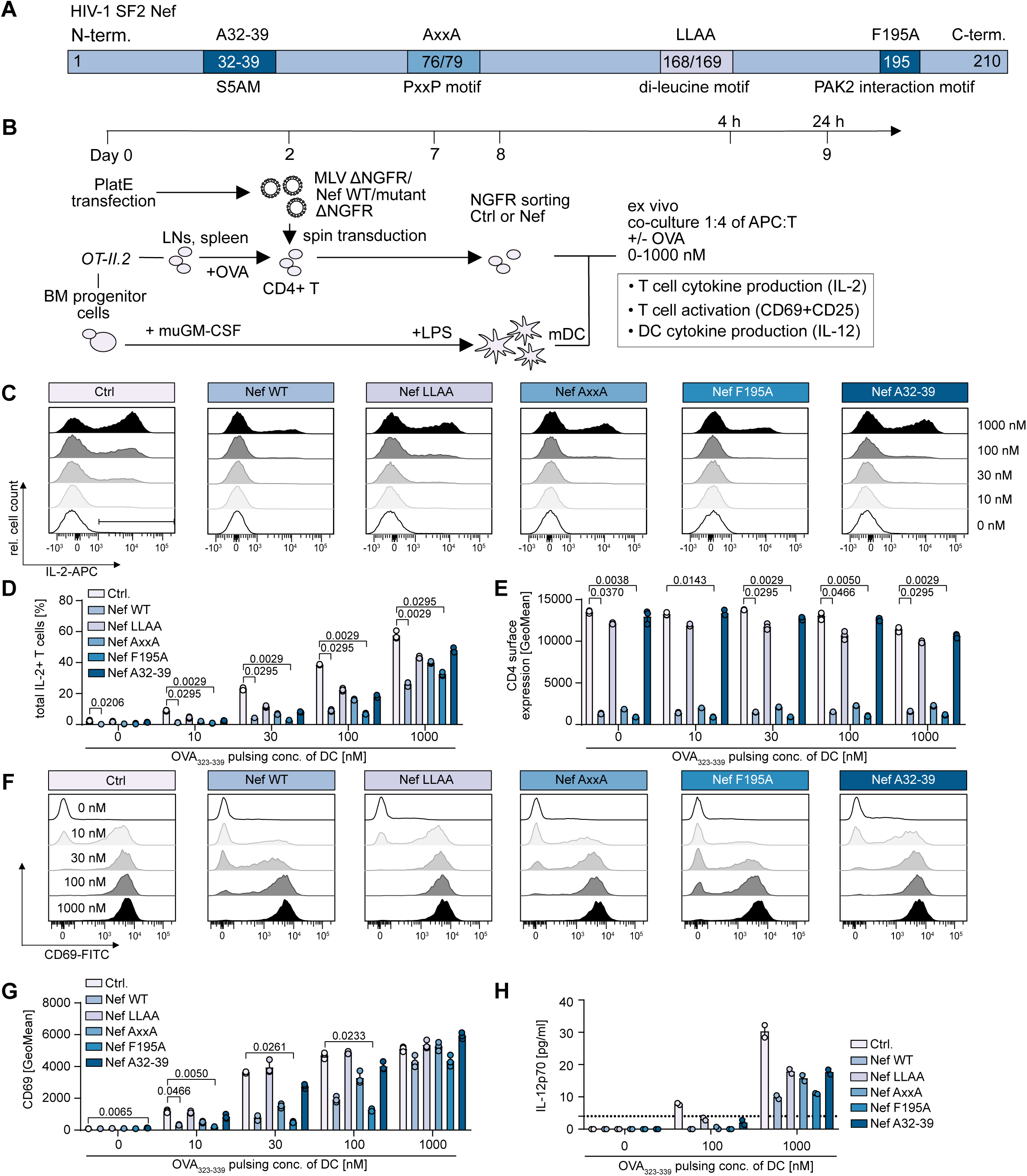
HIV-1 Nef interferes with CD4 T cell-DC interactions via its dileucine and S5AM motif. **(A)** Schematic of the HIV-1 SF2 Nef protein and key molecular motifs. **(B)** Experimental set-up. **(C)** Representative histograms showing intracellular IL-2 levels for Ctrl or Nef/Nef mutant cells for indicated conditions in mDC co-culture. **(D)** Quantification of C. **(E)** Quantification of CD4 surface expression for conditions in D. **(F)** Representative flow cytometry histograms showing CD69 expression on CD4 T cells for conditions as in D and E at 24h. **(G)** Quantification of F for the indicated conditions. **(H)** Quantification of IL-12p70 levels in supernatants of 24h co-cultures of Ctrl or Nef and mutant cells with mDCs for indicated antigen pulsing concentrations. Dashed line indicates the detection limit for the assay. D, E, G and H. Shown are means with SD of biological triplicates, i.e. means of technical duplicates with range for H, data is derived from one representative experiment (two additional repeats in Fig. S8). Statistical analysis in D, E and G was performed using Kruskal-Wallis test. Per condition, SF2 Nef and mutant CD4 T cell conditions were compared with the Ctrl CD4 T cell condition only. p-values are indicated for statistically significant differences.

### CD4 downmodulation is a key mechanism by which HIV-1 Nef interferes with CD4 T cell-DC interactions

CD4 is an important co-receptor that stabilizes TCR-MHC class II interactions to enhance antigen sensitivity of T cells which is reflected by enhanced production of IL-2 and IFNγ upon re-stimulation at low antigen doses (*68–70*) (Fig. 6A). We therefore hypothesized that the deficits in CD4 T cell function upon Ag-specific engagement by DCs induced by HIV-1 Nef may stem from reducing CD4 cell surface exposure. To test such a mechanistic link, we performed highly efficient, activation-neutral gene editing to generate *CD4* knock out (KO) OT-II.2 CD4 T cells for functional characterization of homogenous cell populations without detectable CD4 expression (*71, 72*) (Fig. 6B and Fig. S9A). CD4 KO strongly reduced the ability of Ctrl cells to respond to DC antigen presentation with cytokine production and CD69 upregulation (5.2±0.8% for CD4 KO versus 12.8±0.4% IL-2+IFNγ+ cells for NT cells for 1000 nM co-cultures, respectively and CD69 GeoMean of 395.3±23.5 for CD4 KO cells versus 3203.0±83.2 for NT cells for 30 nM co-cultures) (Fig. 6C-G and Fig. S9B-D and F-H) and abrogated DC cytokine responses (Fig. 6H and Fig. S9E and I). CD4 thus plays an essential role in CD4 T cell-DC interactions in this experimental setting and the effects of CD4 depletion thus mirrored those of Nef expression. Presumably reflecting that Nef expressing T cells still displayed residual levels of CD4 at their surface, the disruption of T cell function by complete depletion of CD4 were more pronounced than those exerted by Nef. Moreover, and in line with the partial disruption of Nef function by the AxxA mutation, which does not affect CD4 downregulation by the viral protein (Fig. 5E), combining Nef expression with CD4 KO had an additive effect on the suppression of cytokine production (1.0±0.1% IL-2+IFNγ+ cells for Nef WT CD4 KO versus 5.2±0.8% for Ctrl CD4 KO and 2.7±0.1% for Nef LLAA CD4 KO in 1000 nM co-cultures). Collectively, these results define that CD4 downmodulation by HIV-1 Nef is essential to impair CD4 T cell-DC communication.

**Fig. 6.**
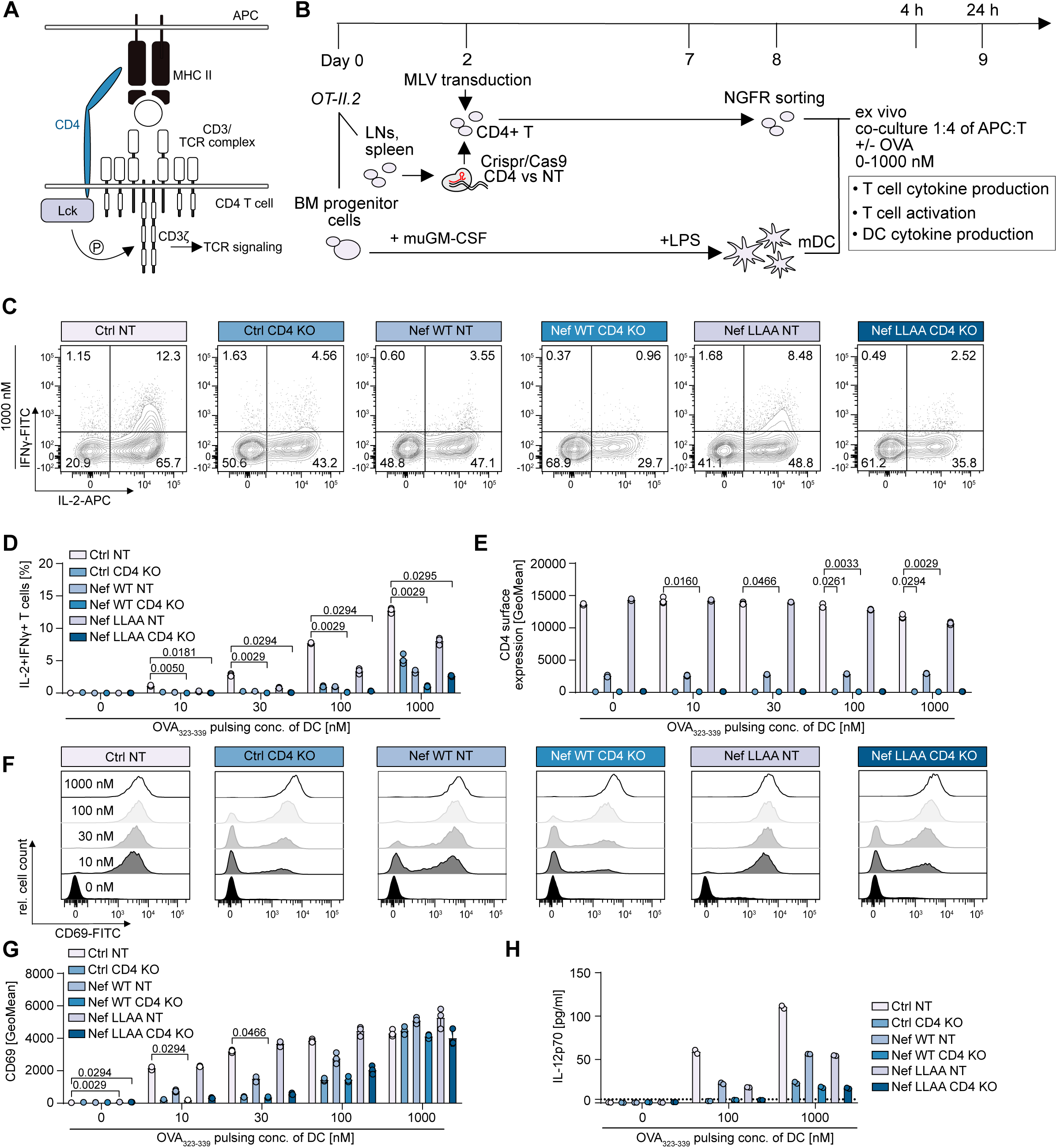
CD4 downmodulation is an essential effector function by which HIV-1 Nef interferes with CD4 T cell interaction with DCs. **(A)** Schematic of the CD4 co-receptor function in the context of TCR-MHC class II interaction. **(B)** Schematic of the experimental workflow. **(C)** Representative flow cytometry contour plots showing IL-2 and IFNγ production at 4.5h of NT or CD4 KO OT-II.2 cells transduced with Ctrl or Nef WT or LLAA mutant MLV in co-culture with mDC pulsed with 1000 nM OVA323-339. **(D)** Quantification of IL-2 and IFNγ-positive CD4 T cells for conditions in C, at respective antigen-pulsing concentrations. **(E)** Quantification of surface CD4 levels for conditions in D. **(F)** Representative flow cytometry histograms showing surface CD69 expression on OT-II.2 cells at 24h in co-culture with mDCs for the indicated conditions. **(G)** Quantification of F. **(H)** Quantification of IL-12p70 levels by ELISA in supernatants of 24h co-cultures of OT-II.2 cells with mDC pulsed with 0, 100 or 1000 nM OVA323-339. Dashed line indicates the detection limit for the assay. D, E, G and H. Shown are means with SD of biological triplicates, i.e. means of technical duplicates with range for H, data is derived from one representative experiment (two additional repeats in Fig. S9). Statistical analysis in D, E and G was performed using Kruskal-Wallis test. Per condition, SF2 Nef and mutant CD4 T cell conditions were compared with the Ctrl CD4 T cell condition only. p-values are indicated for statistically significant differences.

## DISCUSSION

Our results demonstrate that expression of the HIV-1 pathogenesis factor Nef in CD4 T cells profoundly impacts their response to stimulation by DC-presented antigens (Fig. S10). Nef-mediated impairments of CD4 T cell function included reduced activation and Th1 polarization associated with skewed cytokine production profiles. These functions reflected the deregulation of transcriptional responses by reducing expression of genes associated with TCR-signaling and, in particular, Th1-polarization, while enhancing transcription of Nef interaction partner genes that enable TCR-independent activation. Since impaired Th1 polarization persisted even during extensive cell proliferation, Nef imprints a unique transcriptional signature in CD4 T cells that translates in altered cytokine production and leads to inefficient Th1 polarization upon antigen-specific activation. Importantly, our results reveal that Nef not only affects CD4 T cells but also interacting DCs as we observed impaired DC activation on a transcriptional and functional level. This was in particular reflected by a suboptimal induction of co-stimulatory and maturation-associated transcriptional pathways and consequently, reduced release of pro-inflammatory factors specifically associated with Th1 T cell polarization and help to CD8 T cells. Th1 polarization is thus impaired by Nef via direct effects on CD4 T cells as well as indirect effects on DC function, which in turn creates a feedback loop that further impairs Th1 polarization (Fig. S10, bottom panel). These effects of Nef are strikingly similar to observations in PLWH of inefficient activation-induced IL-2 and IFNγ production of both HIV-specific and unspecific CD4 T cells (*11, 12, 73, 74*) as well as the reduction of Th1-associated gene expression and DC maturation and CD28-costimulation pathways upon HIV infection of *ex vivo* tonsil cultures and HIV reservoir cells in PLWH on ART (*75, 76*). Since HIV-specific CD8 T cell responses require CD4 help mediated by CD4 T cell interactions with DCs and CD4 T cell secretion of IL-2 (*77, 78*), these impairments will limit breadth and efficacy of CD8 T cell responses. Expressed to high levels during acute infection, Nef is also synthesized to functional levels from latently infected or low-replicating HIV infected cells even under ART (*79*). In addition, functional Nef protein can be delivered to bystander cells by several mechanisms including secretion in extracellular vesicles and transfer by cell protrusions (*80–86*). The pathological effects of Nef are thus not limited to acutely infected cells, which could explain the persistence of immune dysfunction under ART.

In addition, production of IL-7, which can promote HIV replication and increase production from reservoir cells, was elevated by the presence of Nef and is consistent with Nef’s ability to boost virus replication *in vivo* (*54, 55*). Collectively, these results define the pathogenesis factor Nef as key driver of CD4 T cell dysfunction and impaired Th1 polarization in HIV infection and suggest this mechanism to contribute to generalized CD8 T cell dysfunction while promoting virus spread in PLWH.

Studies of CD4 T cell - DC communication using bacterial superantigen are limited by the fact that superantigen-driven T cell activation bypasses CD4 co-stimulation and Lck signaling by stably connecting TCR and MHC class II (*70, 87*). Consequently, a previous study investigating the effects of HIV-1 Nef on superantigen-primed CD4 T – DC cell interaction with DCs could not detect effects of HIV-1 Nef on T cell-DC interactions (*42*). Here, the use of monoclonal mouse T cells, in which all cardinal Nef functions are preserved (*43, 88, 89*), enabled us to study CD4 T cell responses to a specific peptide antigen and thus to overcome this limitation. With disruption of Th1 polarization and antigen-specific communication between CD4 T cells and DCs, which is key for CD8 T cell function during chronic infection, our results define new effector functions of HIV-1 Nef that likely contribute to its ability to promote immune evasion and accelerate disease progression in PLWH. Our mutagenesis and gene editing analyses identified the downregulation of CD4 from the cell surface of CD4 T cells as a key mechanism by which the multifunctional Nef protein interferes with CD4 T-DC communication. CD4 downregulation is a highly conserved activity of lentiviral Nef proteins. However, Nef variants impaired in CD4 downregulation are enriched in elite controller PLWH that display superior CD8 T cells responses (*90–93*), suggesting a direct link between this Nef function and host control of HIV infection.

Previously, Nef-mediated CD4 downregulation was thought to prevent superinfection (*94*). While this effect can be demonstrated in *in vitro* infection models, evidence for this role *in vivo*, where co-infection of individual cells as a result of cell-associated virus transmission is readily observed (*95–99*), has been lacking. Our identification of an immunological effector function of Nef-mediated CD4 downregulation provides a new rationale for the central role of this activity in AIDS pathogenesis. Experiments with CD4 blocking antibodies and cells from CD4 KO animals established that CD4 stabilizes TCR-MHC class II interactions and recruits Lck to the TCR complex and hence, increases the sensitivity of CD4 T cells to antigen stimulation (*17, 68, 69, 100*). Our results with gene edited cells confirm this role in the context of acute depletion of CD4 from differentiated cells. In this scenario, the transcriptional deregulation of Nef-expressing CD4 T cells is a direct result of their reduced sensitivity to activation imprinted by downregulation of cell surface CD4 by Nef and the prolonged interactions times we observed between Nef-expressing CD4 T cells and DCs *in vivo* likely represent an attempt to compensate for this signaling defect. Our results further reveal that CD4 downregulation is necessary but not sufficient for CD4 T cell reprogramming by Nef, suggesting synergy with the intracellular rewiring of TCR signaling induced by the viral protein mediated by its SH3-interaction motif, e.g. by translocation of the TCR-proximal kinase Lck (*101, 102*).

The discovery that Nef interferes with CD4 T cell help to DCs establishes that the viral protein globally interferes with the helper function of infected CD4 T cells. Importantly, disruption of help to B cells does not involve CD4 downregulation (*34*) and is thus mediated by distinct mechanisms than the disruption of CD4 T cell-DC communication. The need for distinct mechanisms to disrupt antigen-specific communication of CD4 T cells with different types of APCs emphasizes the different architecture and biology of T-B cell and T-DC ISes (Fig. S11A): while T-B cell synapses are very stable, highly organized and require distinct adaptor molecules such as SAPs for efficient signal transduction, T cell-DC synapses are dynamic and short-lived, lack distinct spatial organization, and do not involve SAPs (*103*). The relatively low rigidity of T-DC ISes appears to be compensated by the involvement of CD4 as important co-stimulant and contributior to synapse stability, which appears to be dispensable at the more stable T-B cell IS (Fig. S11B) (*34, 103, 104*). HIV-1 Nef emerges as versatile tool to mechanistically dissect in more detail the molecular mechanisms that regulate CD4 T cell ISes with different APCs. The evolution of tailored mechanisms to specifically interfere with distinct immune cell communication modes, the high level of conservation of this Nef activity, and the existence of potential back-up mechanisms in the CD4 downregulation function of the viral proteins Vpu and Env (*94*) underscore the relevance of these processes to HIV pathogenesis. Nef employs these distinct mechanisms to interfere with the function of different IS types *in vivo* in order to create a microenvironment that disfavors the development of cellular immunity and to systemically disrupt adaptive immune responses. Strategies to therapeutically target the HIV pathogenesis factor Nef will need to interfere with this emerging role as important contributor to global immune cell dysfunction in PLWH.

### Limitations of the study

Our study is based on the isolated expression of the HIV-1 Nef protein in CD4 T cells via retroviral transduction. While this recapitulates the delivery mode and expression levels in HIV infected cells, we cannot exclude that mechanisms exerted by other viral proteins also affect CD4 T cell-DC communication. Such infection experiments were precluded by the requirement of antigen specific primary immune cells, which are available in mouse models that are not permissive for HIV infection but difficult to establish with human cells. Future studies will focus on studying responses of tonsil histocultures to recall antigen explants. However, while our present study in the OVA system focused on one TCR, variations in e.g. affinity and avidity of naturally selected TCRs are expected to influence the outcome of such experiments. Furthermore, we identified the highly conserved CD4 downmodulation function of Nef as essential to disrupt T-DC communication, but more studies are required to characterize the additional mechanism mediated by the Nef SH3 domain interaction motif that is alsoinvolved.

## MATERIALS AND METHODS

### Study Design

This study was designed to investigate whether expression of the HIV-1 Nef protein in CD4 T cells affects antigen-specific interaction with DCs as a key process towards mounting of adaptive immune responses. To this end, Nef was introduced into murine, transgenic CD4 T cells by retroviral transduction, corresponding to the natural delivery mode and expression level and preserving all major Nef functions. DCs were generated from bone marrow, loaded with antigen and interactions of T cells and DCs characterized *ex vivo* and *in vivo*. A combination of multicolour flow cytometry, cytokine analysis by ELISA, scRNAseq as well as intravital 2-photon microscopy was then used to assess quality of the interactions in an antigen dose-dependent and time-resolved manner. Lastly, a mutagenesis approach, combined with CRISPR/Cas9 KO, was used to mechanistically dissect Nef’s mechanism of action.

### Cell Culture

Plat-E cells were cultured in DMEM high Glutamax containing 1% Penicillin/Streptomycin as well as 1 µg/ml Puromycin and 10 µg/ml Blasticidin for selection of MLV Gag-Pol and Env eco gene expression, respectively (*105*). Cells were split every 2-3 days.

### Mice

All mice used in this study were held and bred at the animal facility (IBF) at Heidelberg University, Heidelberg, were male, between 7-15 weeks of age and of C57BL/6N origin. CD4 T cell donor mice expressed either the transgenic OT-II receptor (Y-chromosome linked inheritance) that recognizes the chicken ovalbumin protein peptide OVA323-339 (OT-II.2 mice) or transgenic, LCMV GP61-80 specific TCR (SMARTA mice) in the context of MHC class II I-Ab expression. SMARTA mice were bred to express the CD45.1 isoform of the CD45 receptor while all other mice were bred to express the CD45.2 isoform to be able to distinguish adoptively transferred cells by antibody staining upon flow cytometry. Transgenic, fluorescent mouse lines serving as donor animals for imaging studies (OT-II.2cfp and P14-mT or OT-II.2_mT/mG mice) were created by crossing transgenic mouse lines with CAG-eCFP, mT/mG mice (*34*). For 2-Photon microscopy experiments, recipient animals were purchased from Janvier Labs (Le Genest-Saint-Isle, France) and were transferred to the BSL-3 animal housing unit of the Center for Integrative Infectious Disease Research of Heidelberg University. Animal projects were registered under T-39/19, T-25/22 and G-300/19 and were approved by the state of Baden-Wuerttemberg, Germany.

### Production of MLV vector

Plat-E cells were seeded without selection antibiotics and were transfected on the following day using following components for transfection of 1×15 cm dish

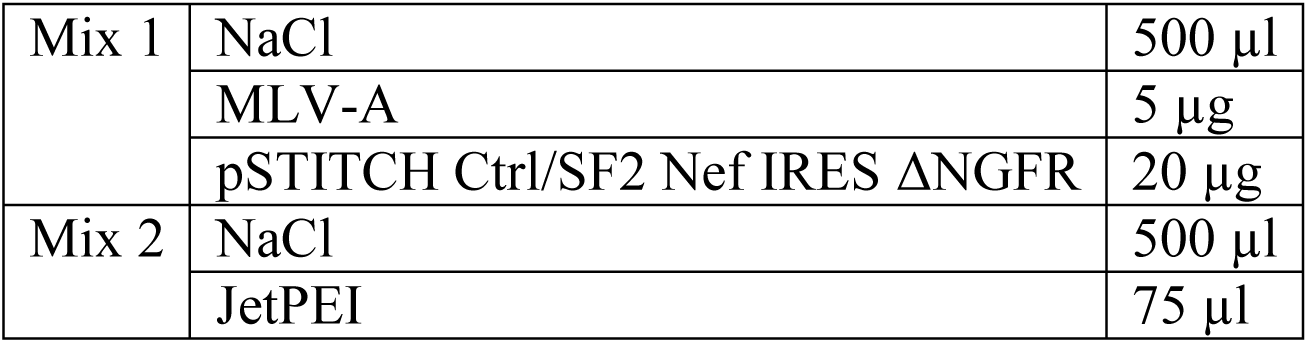

|  |  |  |
| --- | --- | --- |
| Mix 1 | NaCl | 500 µl |
|  | MLV-A | 5 µg |
|  | pSTITCH Ctrl/SF2 Nef IRES ΔNGFR | 20 µg |
| Mix 2 | NaCl | 500 µl |
|  | JetPEI | 75 µl |

Mix 1 and 2 were prepared separately, then mixed together and incubated for 20 min at RT and added to Plat-E dishes. Medium was changed 6h later without addition of selection antibiotics and MLV supernatant harvested 2 days later and filtered through a 0.45 µm filter to remove cell debris. pSTITCH-GFP and MLV-A constructs were a kind gift from Reno Debets (Department of Medical Oncology, Erasmus MC, Rotterdam, The Netherlands) and Nef alleles as well as NGFR reporter were inserted as described previously (*34, 43*).

### Isolation and activation of murine lymphocytes

For *ex vivo* co-culture studies, *in vivo* activation and proliferation and intravital microscopy analysis, lymphocytes from OT-II.2 or OT-II.2cfp mice were used as indicated. In short, inguinal, popliteal, axillary, brachial and cranial lymph nodes and spleen were harvested and passed through a 70 µm cell strainer, rinsed and washed once prior to ACK lysis for 5 min at RT. Cells were washed once, counted and cultured at 5E06 cells/ml in CMR with the OVA323-339 peptide at 1 µg/ml concentration. Cells were incubated for 2 days with peptide and then used for transduction with MLV.

### Transduction of murine lymphocytes

24-well plates were pre-treated with Retronectin (working concentration of 8-12 µg/ml) by adding 0.5 ml Retronectin solution per well, incubating the plates for 3h at 37°C followed by washing 2x with 1xPBS. Per well, 2.5E06 CD4 T cells were added to 2 ml of MLV supernatant, spin transduced at 37°C for 2.5h at 2.300 rpm and incubated for another 4-6h after which cells were collected from the wells and resuspended at 1-2E06 cells/ml in a total volume consisting of 1/3 of MLV supernatant and 2/3 fresh CMR supplemented with murine IL-7 at 10 ng/ml. Cells were then incubated for another 5-6 days with renewal of 1/2 to 2/3 of the medium with fresh CMR and supplementation of fresh IL-7 every 2-3 days.

### NGFR-sorting of transduced murine lymphocytes

On day 5-6 post transduction, murine lymphocyte cultures were collected, counted and subjected to NGFR-sorting based on NGFR-reporter expression of transduced cells using EasySep™ Human CD271 Positive Selection Kit II (Stemcell Technologies) according to the manufacturer’s instructions with three consecutive purification steps. T cells were then either used directly or stained with Celltrace violet or CFSE for proliferation studies. T cell concentration was then adjusted to 2E07/ml equaling 3E06 cells in 150 µl for retro orbital injection into recipient mice for *in vivo* experiments or to 1-2E06 cells/ml for *ex vivo* co-culture with DCs.

### BMDC generation from murine bone marrow

For *ex vivo* co-culture studies and *in vivo* activation and proliferation analysis, BMDCs were generated from the same animals used for lymphocyte isolation while for intravital 2-photon microcopy, mT-reporter mice (P14-mT or OT-II.2_mT/mG) were used based on a method described by Lutz et al. (*106*). Briefly, hind legs were removed from euthanized mice, bones were extracted and cleaned using lint-free paper towels and disinfected by short immersion in 70% ethanol. Bone ends were then opened with scissors and flushed with PBS in a 10 ml syringe with a 27G needle to extract bone marrow. Bone ends were then further disintegrated with scissors in PBS and, together with bone marrow, passed over a 100 µm cell strainer to remove bone fragments. Cells were centrifuged for 5 min at 1200 rpm, washed with RPMI 1640 without supplements and resuspended in BMDC medium. Cells were counted, omitting very small cells (erythrocytes and platelets) and 3E06 cells were cultured in 10 cm petri dishes in 10 ml BMDC medium supplemented with 20 ng/ml recombinant murine GM-CSF. On day 3 of culture, another 10 ml BMDC medium with fresh GM-CSF were added and on day 6, half of the medium was carefully removed and replaced with 10 ml BMDC medium with fresh GM-CSF. BMDCs were then either incubated until day 8 or treated with 1 µg/ml LPS on day 7 to induce maturation.

### *Ex vivo* DC pulsing with peptides

BMDCs were diluted at 5E06 cells/ml in BMDC medium and OVA323-339 added at 10, 30, 100 or 1000 nM concentration. Cells were incubated for 45 min at 37°C, washed at least 2x with BMDC medium and cell concentration was adjusted for injection into mice at 5E07/ml (5E05 cells in 10 µl per injection) or *ex vivo* co-culture with T cells at 0.25-0.5E06 cells/ml.

### *Ex vivo* co-culture of murine CD4 T cells and BMDC

BMDCs and T cells were co-cultured at a ratio of 1:4 in 96-well plates with total volumes of 100-200 µl/well. Biological triplicates were prepared per condition. For activation analysis, cells were incubated for 24h and harvested while proliferation was assessed at 72h. For cytokine analysis, cells were co-cultured for 0.5h and Monensin was added to the cultures for the remaining 4h. For cytokine analysis by ELISA, supernatants of triplicate wells were pooled at 24h and analyzed in technical duplicates.

### *In vivo* activation and proliferation analysis

Recipient mice for activation and proliferation analysis were SMARTA CD45.1 (transgenic or non-transgenic for the SMARTA T cell receptor) for OT-II.2 (CD45.2) CD4 T cell transfer. In total 3E06 T cells were injected in a total volume of 150 µl of NaCl retroorbitally into recipient mice under short-term isoflurane anesthesia. One day later, mature BMDC were injected subcutaneously above the ankle of all four legs of recipient mice at 5E05 cells in 10 µl total volume per injection under isoflurane anaesthesia. Each extremity received BMDC with a different peptide-pulsing concentration (0 and 10 nM for front legs to 30 and 100 nM for hind legs). Approximately 16h post BMDC transfer, FTY720 was injected intraperitoneally at 1 mg/kg concentration in 150 µl of total volume to block lymphocyte egress from lymph nodes and 24h or 72h post FTY720 treatment, mice were euthanized by cervical dislocation for activation and proliferation analysis. From each mouse, draining lymph nodes of BMDC injection sites were harvested. For front legs, axillary and brachial lymph nodes were harvested for each side while popliteal lymph nodes were harvested for the hind legs. Lymph nodes were transferred into culture medium containing collagenase/dispase at 1 mg/ml, mechanically dissociated and incubated for 30 min at 37°C on a rocking incubator. Cells were then passed over a 70 µm cell strainer, washed with FACS buffer and resuspended in antibody staining solution for detection of CD3-APC-Cy7, CD45.2-PE and CD69-FITC (all 1:100 diluted in FACS buffer), washed, fixed in 3%PFA and acquired on a BD FACSymphony™ A3 flow cytometer. FlowJo™ v10.7.1 (BD Biosciences) was used for data processing. Values for CD69 (GeoMean) or CTV (Mean) signal on Ctrl or Nef CD45.2 cells were normalized to the Ctrl 100 nM OVA condition for CD69 or the 0 nM OVA condition for CTV analysis per experiment. Percentage of CD45.2 cells among CD3+ cells at 24 or 72h was normalized to the mean of Ctrl T cells in the 100 nM OVA condition per experiment.

### Intravital 2-Photon Microscopy

T cells and mDCs were injected as described above, using cells from OT-II.2cfp and mT-reporter mice and injecting pulsed (1000 nM OVA323-339) or unpulsed mT-DCs only into the hock of the right hind leg, respectively. 20-28h or 36-44h post DC transfer (∼4-12 and 20-28h of encounter with T cells), the right popliteal lymph node of the right hind leg was exposed microsurgically for intravital imaging, as previously described (*43*). Mice were anesthetized with isoflurane (3% for induction and 1-1.5% for maintenance, Isofluran CP, CP-pharma #1214), under Buprenorphine (0.05–0.1 mg/kg, Buprenorphinhydrochloride/Temgesic, 1A Pharma) analgesia and approximately 15µg Meca79-AlexaFluor 647 antibody in 0.9% NaCl was injected intravenously prior to imaging, to label high endothelial venules. Intravital 2-Photon microscopy was performed with a Nikon Eclipse FN-1 upright microscope equipped with a 25× Nikon CFI-Apo (NA 1.1) objective and a TrimScope II 2PM system controlled by ImSpector software (LaVision BioTec), combined with an automated system for real-time correction of tissue drift (*107*). For 2-photon excitation, a Ti:sapphire laser with an optical parametric oscillator (OPO, Coherent MPX Package) were tuned to 840 and 1,100 nm, respectively. For 4-dimensional analysis of cell migration, 16 x–y sections with z-spacing of 4 μm (60 μm depth) were acquired every 20 s for 40 min (121 time steps); the field of view was 300 × 300 μm. Emitted light and second harmonic signals were detected through 480/30-nm, 525/50-nm, 595/50-nm, and 690/50-nm bandpass filters using non-descanned detectors. Timelapse movies were reconstructed using Arivis Vision 4D software version 4.1.2 (Arivis AG). Cell interactions were manually tracked and quantified using VisionVR software version 4.0.1 (Arivis AG), and T cells were semiautomatically tracked and manually controlled using Imaris 10.2.0 (Bitplane). Meca79-AF647, was custom-made using the respective Hybridoma cell line, by the company Nanotools and coupled to AlexaFluor 647 using the Antibody labeling kit AlexaFluor 647 NHS-Ester or Meca79-AF647 from SantaCruz Biotechnology. Of note, antigen-specific interactions with DCs observed in our set-up were in general shorter compared to data from literature due to the memory phenotype of the transduced T cells (*108*).

### Cell preparation for flow cytometry analysis of *ex vivo* co-cultures

Cells from *ex vivo* co-cultures were washed with 1xPBS, and stained with zombie violet cell viability dye according to the manufacturer’s instructions. Cells were blocked with mouse Fc block (1:5000) diluted in FACS buffer for 5 min at RT and resuspended in antibody diluted in FACS buffer for cell surface staining for 20 min at 4°C. Cell surface antibodies NGFR-PE, CD11c-APC, CD25-PerCP-Cy5.5, CD69-FITC, CD69-BV650, CD86-PerCP and CD4-APC-Cy7, combined as indicated, were all diluted 1:100. Cells were then washed in 1xPBS and fixed in 3%PFA for 10 min at RT, washed with 1xPBS and either acquired directly on a BD FACSCelesta™ flow cytometer or stored for up to 1 week at 4°C until acquisition or prior to intracellular staining. FlowJo™ v10.7.1 (BD Biosciences) was used for data processing.

### Intracellular cytokine analysis

Intracellular cytokine staining was performed using BD Cytofix/Cytoperm™ Fixation/Permeabilization Kit (BD Biosciences) according to the manufacturer’s instructions. Briefly, after viability and cell surface staining, cells were pelleted, resuspended in Cytofix/Cytoperm solution and incubated for 20 min at 4°C. Cells were then washed 2x with 1xCytoperm buffer, resuspended in antibody diluted in Cytoperm buffer and incubated for 30 min at 4°C. Both IFNγ-FITC and IL-2-APC antibodies were used at 1:100 dilutions. Cells were then washed with 1xPBS and resuspended in 1xPBS for flow cytometry analysis on BD FACSCelesta™ on the same day. FlowJo™ v10.7.1 (BD Biosciences) was used for data processing.

### Intranuclear transcription factor analysis

Intracellular and intranuclear T-bet expression was assessed using the True-Nuclear™ Transcription Factor Buffer set (BioLegend) according to the manufacturer’s instructions. Briefly, after viability and cell surface staining, cells were fixed with 1xFix working solution for 45-60 min at RT, washed 2x with 1xPerm buffer and incubated with T-bet-PerCP antibody at 1:100 dilution in 1xPerm buffer for 45 min at RT. Cells were washed 2x with 1xPerm buffer and acquired on the same day on BD FACSCelesta™. FlowJo™ v10.7.1 (BD Biosciences) was used for data processing.

### IL-12p70 ELISA

IL12p70 ELISA was performed using the ELISA MAX™ Deluxe Set Mouse IL-12 (p70) from BioLegend according to the manufacturer’s instructions. In short, 96-well NUNC™ plates were coated overnight at 4°C with IL-12p70 capture antibody, blocked for 1h at RT and incubated for 2h at RT with standard or supernatants of *ex vivo* co-cultures that were either used undiluted or diluted 1:2 with assay diluent. Each sample was analyzed in duplicate. Samples were then incubated with Biotin-conjugated detection antibody for 1h at RT prior to addition of Avidin-HRP and incubation for 30 min at RT. TMB substrate was then added and wells incubated for 15 min at RT and reaction stopped by adding 2N H2SO4. Between each of the steps up to TMB substrate addition, plates were washed four times with PBS-T with 1 min soaking time per wash for the final wash before substrate addition. Absorbance was measured using Infinite® 200 Pro microplate reader (Tecan Group) immediately after H2SO4 addition at 450 nm with a reference absorbance at 570 nm that was subtracted from the values afterwards. GraphPad Prism was then used to interpolate sample concentrations based on the manufacturer’s instructions using a sigmoidal, 4-PL curve fitting.

### Multiplex cytokine analysis in *ex vivo* co-culture supernatants

Supernatants from three independent *ex vivo* co-culture experiments of OT-II.2 CD4 T cells and BMDCs were sent for cytokine analysis by Eve technologies (Calgary, AB, Canada) using the Mouse Cytokine 32-Plex Discovery Assay. Data was processed in Microsoft Excel and either displayed as raw concentrations in pg/ml or as log2-transformed values of ratios relative to the control T cell condition with iDC without antigen or as relative values for Nef conditions in % normalized to the respective Ctrl conditions with antigen set to 100% as indicated.

### SgRNA Design, RNP preparation and Crispr/Cas9 RNP nucleofection

SgRNAs were designed using the Synthego Crispr Online Design Tool (Synthego Corporation, Redwood City, CA, USA) and checked individually for suitable combination. 3 sgRNAs were designed for murine CD4 while the non-targeting sgRNA sequence was provided by Synthego. RNPs for each sgRNA were prepared by combining the respective sgRNA (100 µM) and Cas9 (62 µM) in PBS on ice, using 10 µl of sgRNA and 6.45 µl of Cas9 per 50 µl complex volume. Complexes were incubated for 15-20 min at RT and frozen at −80°C until usage. Immediately before nucleofection, RNPs were thawed on ice and the three RNPs for CD4 were combined in equal amounts to a final volume of 24 µl per 1E07 cells. Cells were prepared for nucleofection by washing in PBS and resuspension in 100 µl of nucleofection solution (Lonza P3 Primary Cell Nucleofector™ X kit L) per 1E07 cells and combined with the RNP mix. Cells were then transferred into single nucleofection cuvettes and pulsed with the DN-100 program using the Lonza 4D nucleofector™. Immediately after nucleofection, RPMI 1640 without supplements was added to obtain a concentration of 5E07 cells/ml and cells were incubated for 15-20 min at 37°C to recover. Complete cell culture medium was then added to obtain 2.5E07 cells/ml of which 100 µl were added directly to wells of a 24-well plate containing MLV supernatant for spin transduction.

### Single-cell RNA Sequencing and Analysis

Single-cell analysis was carried out using Evercode™ split-pool combinatorial barcoding by Parse Biosciences. Briefly, transduced and sorted OT-II.2 CD4 T cells and immature or matured BMDCs were co-cultured for 4 or 24h and harvested for single cell analysis. First, co-cultures were pooled per condition and purified over OptiPrep™ density cushion by centrifugation at 600xg for 15 min to obtain highly viable cell populations. Cells were counted and viability was assessed by transmitted light microscopy using Trypan blue staining and analysis of zombie violet staining by flow cytometry, demonstrating viabilities of over 95% of all samples. Per condition, 1E06 cells were prepared for fixation and processed for storage using the Evercode™ Cell Fixation v3 kit from Parse Biosciences according to the manufacturer’s instructions and filtering the cells twice over 40 µm cell strainers, respectively. Fixed and permeabilized cells were stored in aliquots at −80°C for approximately one and a half months and thawed for downstream processing with Evercode™ WT chemistry version 3 kit according to the manufacturer’s instructions. Cells were split into 8 sublibraries and sequenced on Illumina NovaSeq 6K 200 S1 instrument with a sequencing depth of 20.000 reads/cell with paired end sequencing, using two sequencing lanes. FASTQ files were concatenated per sublibrary for R1 and R2 reads, respectively and concatenated files were uploaded and processed using Trailmaker™ pipeline module (https://app.trailmaker.parsebiosciences.com/; pipeline v1.4.1, Parse Biosciences). Firstly, unfiltered count matrices were uploaded and cells were filtered for a minimum number of transcripts per cell on a per sample basis (min. threshold range 122.94 to 765.34). At this step, the set of iDC co-cultures with Ctrl or Nef T cells at 4h (with and without antigen, well IDs A2, A3, A7 and A8) were filtered out since one sample of this set had insufficient cell counts that prevented further analysis. This was due to a technical problem rather than biological variation. Next, a mitochondrial content filter was applied to remove dead or dying cells from further analysis, excluding barcodes with high mitochondrial content (max. 4.25-16.16% cut-off). This was followed by exclusion of outliers that deviated from a spline regression model between number of genes vs number of transcripts (p-values between 0.000133 and 0.001). Lastly, cells with a high probability of being a doublet were filtered out by applying the scDblFinder method (probability threshold between 0.38692 to 0.76466). Data normalization (LogNormalize), principal-component analysis (PCA, based on 2000 highly-variable genes, 26 PCs that explain 90.01% of variation) and data integration were performed using Harmony (Seurat analysis tool) on high quality cells. Unifold Manifold Approximation and Projection (UMAP) embedding was calculated for visualization with a minimum distance of 0.3 using cosine distance metric. Clusters were further identified by the Louvain method with 0.6 resolution. Clusters were annotated based on their marker gene expression and associated pathways (top 50 genes entered for Enrichr pathway analysis, (*109–111*)) that distinguished the cells from all other cells by Wilcoxon rank-sum test. Total T cells and total DCs, respectively, were further subgated and subset into new projects for celltype-specific data integration and embedding that allowed for more detailed analysis. Notably, we identified a subset of DC clusters associated with T cell signaling signatures (‘T cell-like’ and ‘TCR-signaling’ subclusters) that expressed *Cd247* and several other T cell-associated genes as well as DC marker genes such as *Cd74*, *Fscn1* and others (Fig. 4C, Fig. S4B, Fig. S5W). As examination of read counts in those clusters did not indicate the population represents doublets of T cells and DCs and the cells participated in the general DC response to antigen-specific interaction with T cells by gene upregulation in a similar manner to other DC clusters we did not exclude these cells from analysis of total DC responses. Analysis of relative cluster contributions and differential gene expression was performed with integrated tools in the Trailmaker™ software. For differential gene expression analysis between groups of samples, volcano plots were generated and significance calculated by inbuilt pseudobulk limma-voom workflow in Trailmaker™. For analysis of specific antigen-induced genes (volcano plots), samples of Ctrl and Nef T cells in co-culture with mDC (4+24h timepoints) and iDC (24h) were pooled for – Ag versus +Ag conditions, respectively. For evaluation of Ctrl versus Nef conditions at indicated timepoints for T cells or DCs, individual samples were compared for 4h timepoints (mDC co-cultures) or pooled for 4+24h timepoints (mDC and iDC co-cultures). For pathway enrichment analysis, Ctrl and Nef samples (one sample per condition) were first compared using the inbuilt pseudobulk limma-voom workflow that provided a table of average expression and logFC values for sample versus sample comparisons. Genes with positive or negative logFC were then separately ranked according to average expression and magnitude of the logFC and a combined rank was calculated to obtain a list of the top 100 genes with the highest average expression and highest magnitude of logFC. These top 100 genes were then used for pathway enrichment analysis using MSigDB (Mouse MSigDB v2024.1.Mm updated August 2024, (*112–114*)). Pathway analysis was performed with the Reactome 2024 database.

For sequencing data deposition, concatenated FASTQ files were split per sublibrary according to well ID of the initial barcoding round to exclude samples not related to this study. For this purpose, a python code was provided by Parse Biosciences and adapted (https://support.parsebiosciences.com/hc/en-us/articles/29780938849044-Partitioning-fastq-files-by-round-1-well-coordinates).

### Statistical analysis

Data analysis, visualization and statistical analysis of all datasets, unless otherwise indicated, was carried out using Microsoft Excel and Prism version 8.0.1 (GraphPad). Normality was tested for each dataset using Shapiro-Wilk test and statistical significance was calculated using the appropriate parametric, i.e. non-parametric tests as indicated in the Figure legends. For statistical analysis of scRNAseq data, inbuilt analyses in Trailmaker™ (Parse Biosciences) and/or inbuilt analyses of MSigDb or Enrichr were used for pathway enrichment analysis as detailed in the respective section.

## Supporting information

all supplementary material

## Supplementary Materials

Figs. S1 to S11.

Table S1 and S2.

Data S1

Movies S1 to S6

## Acknowledgments

We would like to thank the Interfaculty Biomedical Research Center (IBF) at Heidelberg University for support with mouse breeding and caretaking. CAG-eCFP and mT/mG mice were kind gifts from Anette Oxenius and Marc Freichel, respectively, pSTITCH and MLV-A plasmids were kindly provided by Reno Debets. We are grateful for microscopy support from the Infectious Diseases Imaging Platform (IDIP) at the Center for Integrative Infectious Disease Research, Heidelberg and the FACS Core Facility (dFCCU) of the Medical Clinic, Heidelberg University Hospital. The authors gratefully acknowledge the data storage service SDS@hd supported by the Ministry of Science, Research and the Arts Baden-Württemberg (MWK) and the German Research Foundation (DFG) through grant INST 35/1503-1 FUGG. Lastly, we want to thank the Next Generation Sequencing Core Facility at the German Cancer Research Center (DKFZ), Heidelberg. OTF and MP are members of the SynthImmune cluster of excellence (EXC3018-1).

## Funding

Deutsche Forschungsgemeinschaft (DFG, German Research Foundation), Projektnummer 240245660 - SFB 1129 (project 8)

Ministry of Science, Research and Arts Baden-Württemberg, Germany (support during the establishment phase of the SynthImmune cluster of excellence)

German Center of Infection Research (DZIF, project immune control, TTU HIV)

The Health and Life Science Alliance Heidelberg-Mannheim funded by the Ministry of Science, Research and Arts Baden-Württemberg, Germany

## Author Contributions

Conceptualization: OTF, BS and KMo

Methodology: KMo, BS, KMa and JR

Investigation: KMo and BS

Resources: KMo, BS, JR and MP

Data analysis: KMo, JR and BS

Funding acquisition: OTF and MP

Supervision: OTF

Writing—original draft: KMo and OTF

Writing—review and editing: KMo, BS, MP and OTF

## Data and Materials availability

This study did not generate new unique reagents or report original code. All primary data generated in this study necessary for evaluation of conclusions are presented in the main and supplementary figures. ScRNAseq data has been submitted to Gene expression omnibus database (GEO, NCBI) and will be available with accession number (accession number will be submitted as soon as possible). All relevant materials and antibodies used for flow cytometry are included in the supplementary table S2. Any additional information required to reanalyze the data reported in this paper is available from the Lead Contact upon request.

