## Supplementary material for "HIV-1 Reprograms CD4 T Cell Responses by Impairing Antigen-specific Communication with Dendritic Cells": all supplementary material

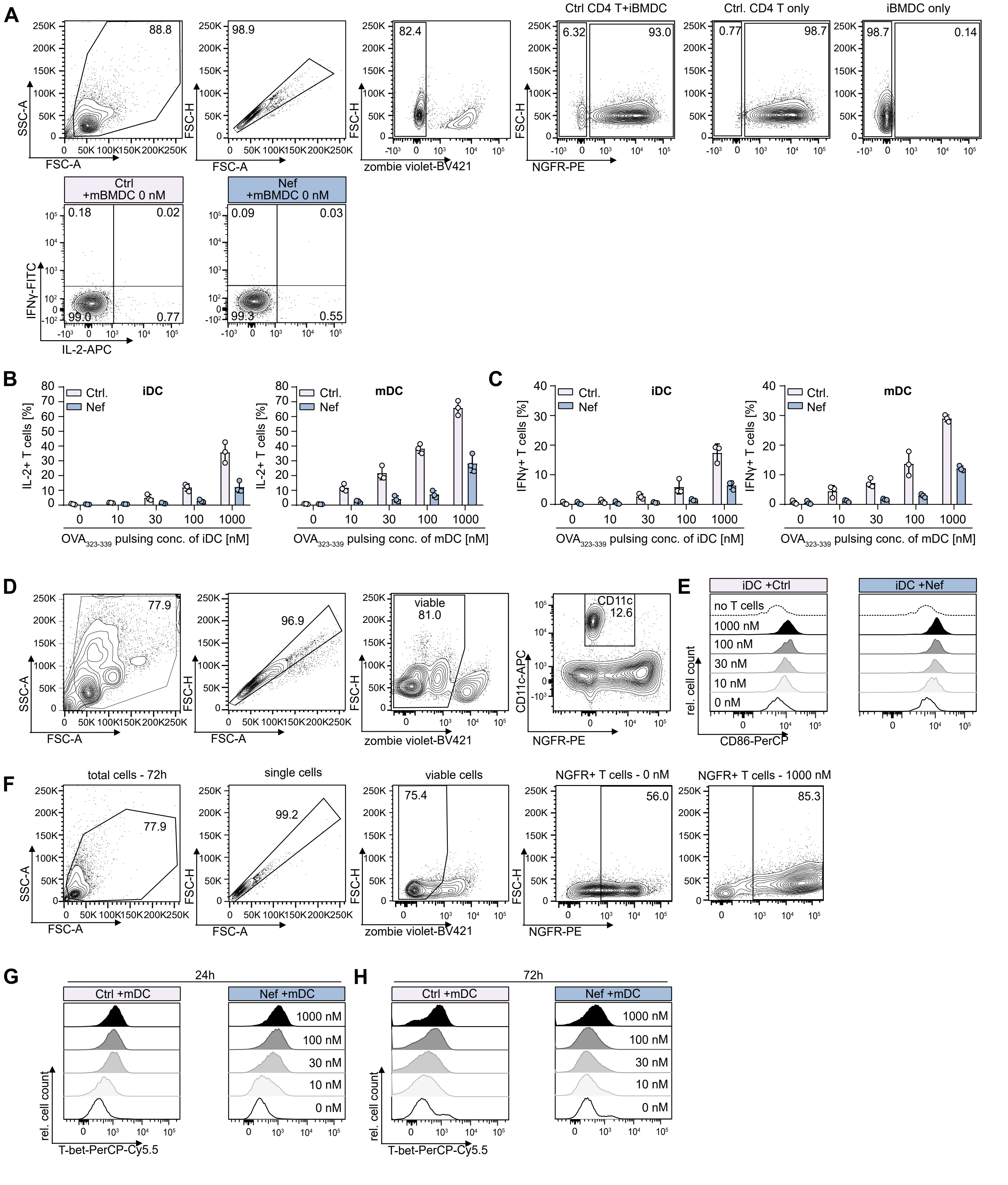

A
SSC-A
FSC-A
88.8
FSC-H
98.9
82.4
zombie violet-BV421
Ctrl CD4 T+iBMDC
6.32
93.0
NGFR-PE
Ctrl. CD4 T only
0.77
98.7
iBMDC only
98.7
0.14
Ctrl
+mBMDC 0 nM
0.18
0.02
99.0
0.77
IFNγ-FITC
IL-2-APC
Nef
+mBMDC 0 nM
0.09
0.03
99.3
0.55
B
Ctrl.
Nef
iDC
mDC
IL-2+ T cells [%]
OVA323-339 pulsing conc. of iDC [nM]
OVA323-339 pulsing conc. of mDC [nM]
C
IFNγ+ T cells [%]
D
77.9
96.9
viable
81.0
CD11c
12.6
CD11c-APC
E
iDC +Ctrl
iDC +Nef
no T cells
1000 nM
100 nM
30 nM
10 nM
0 nM
rel. cell count
CD86-PerCP
F
total cells - 72h
77.9
single cells
99.2
viable cells
75.4
NGFR+ T cells - 0 nM
56.0
NGFR+ T cells - 1000 nM
85.3
G
24h
Ctrl +mDC
Nef +mDC
T-bet-PerCP-Cy5.5
H
72h

**Fig. S1. HIV-1 Nef impairs CD4 T cell cytokine production and Th1 differentiation but not DC maturation upon cognate interaction *ex vivo*. (A)** Upper panel shows gating strategy to identify OT-II.2 CD4 T cells in DC co-cultures with examples of T cell only/DC only conditions, lower panel shows example for IL-2 and IFNγ signal among subgated NGFR+ cells in co-cultures without antigen. **(B)** and **(C)** Quantification of total IL-2 and IFNγ positive CD4 T cells for indicated conditions. Shown are means with SD of three independent experiments. Each symbol represents the mean of biological triplicates within one independent experiment. Statistical analysis was performed using Wilcoxon test. p-values are indicated for significant differences. **(D)** Gating strategy to identify CD11c+ DCs in co-cultures with CD4 T cells for CD86 analysis. **(E)** Representative histograms showing CD86 expression on iDCs in co-culture with Ctrl or Nef CD4 T cells as indicated. Histogram for the no T cell condition (iDC only) is identical for Ctrl and Nef plots. **(F)** Gating strategy for analysis of intracellular/intranuclear T-bet levels. Cells were gated for singlet signal and viability based on exclusion of zombie violet dye and subgated for NGFR expression. Shown are examples for Ctrl CD4 T cells at 72h without (0 nM) or with (1000 nM) peptide pulsing of DCs. **(G)** and **(H)** Representative histograms showing T-bet expression of Ctrl or Nef CD4 T cells in co-culture with mDCs at 24 (G) or 72h (H), respectively, at indicated peptide pulsing concentrations.

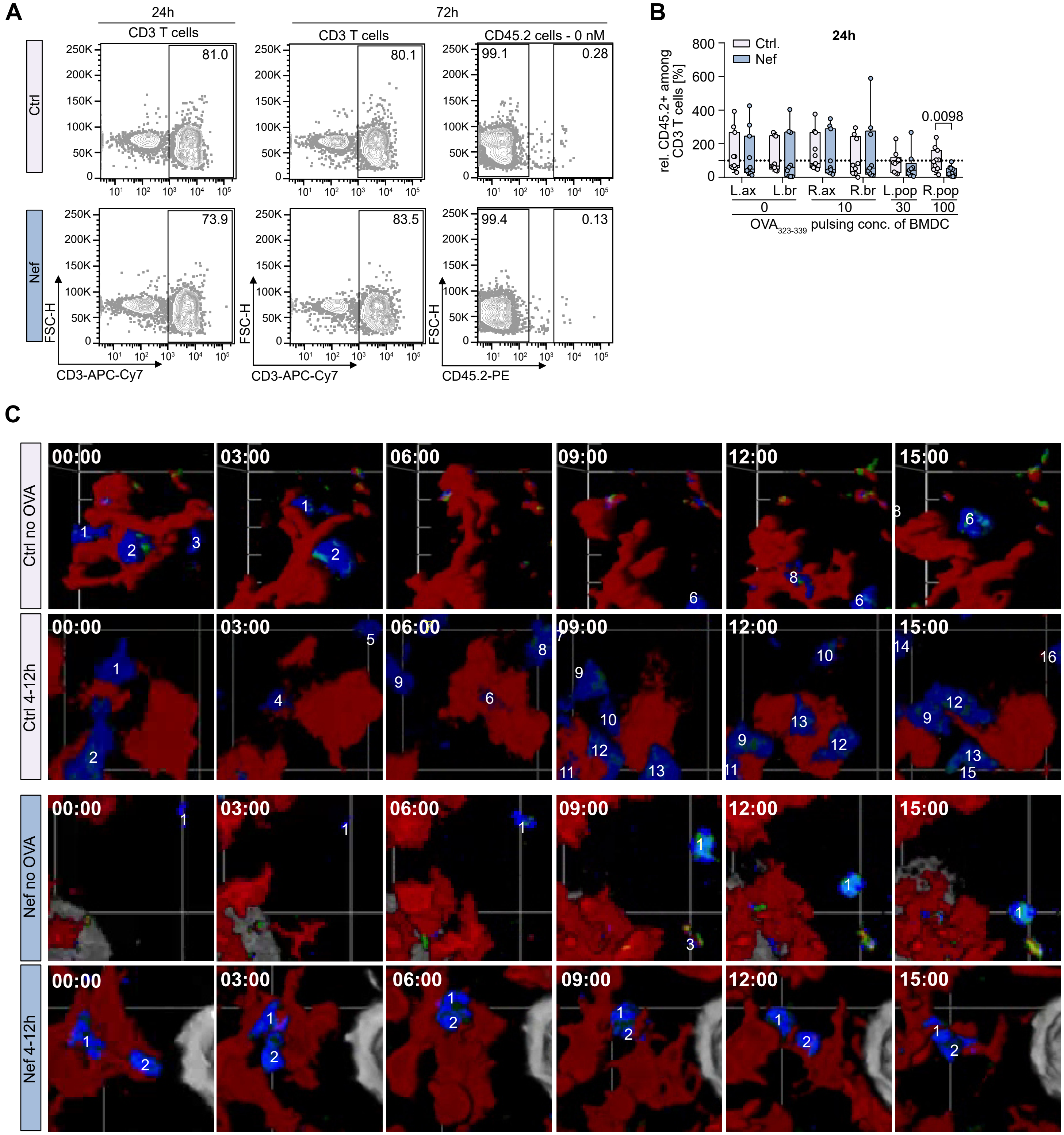
A
24h
72h
CD3 T cells
CD3 T cells
CD45.2 cells - 0 nM
Ctrl
Nef
81.0
80.1
99.1
0.28
73.9
83.5
99.4
0.13
FSC-H
CD3-APC-Cy7
CD3-APC-Cy7
CD45.2-PE
B
24h
Ctrl.
Nef
rel. CD45.2+ among CD3 T cells [%]
0.0098
L.ax
L.br
R.ax
R.br
L.pop
R.pop
0
10
30
100
OVA323-339 pulsing conc. of BMDC
C
Ctrl no OVA
Ctrl 4-12h
Nef no OVA
Nef 4-12h
00:00
03:00
06:00
09:00
12:00
15:00

**Fig. S2. HIV-1 Nef prolongs early antigen-specific interaction dynamics of CD4 T cells and DCs within draining lymph nodes *in vivo*. (A)** Representative flow cytometry plots showing gating of CD3+ cells among total cells obtained from draining lymph nodes of recipient animals 24h (left column) or 72h (middle column) after T cell and BMDC transfer for Ctrl (upper panel) or Nef (lower panel) conditions. Right column displays percentage of CD45.2 cells in draining lymph nodes at 72h without antigen pulsing of BMDCs. **(B)** Frequency of CD45.2 OT-II.2 cells among CD3+ cells at the 24h timepoint normalized to the mean of Ctrl T cells in the 100 nM OVA condition per experiment. Shown is a box and whisker plot with median, min and max range for n=4 individual experiments with 2-3 animals per group per experiment. Each dot represents data from one animal per condition. Statistical analysis was performed using Mann-Whitney test. p-values are indicated where significant differences between Ctrl and Nef conditions were identified. **(C)** Representative micrographs/ sequential still images from intravital 2-Photon microscopy movies (Movies S1, S3, S4 and S6) with timestamps showing sequential T cell-DC interactions for Ctrl (upper panels) or Nef (lower panels) T cells with OVA-pulsed DCs at 4-12h or without antigen as indicated. Numbers mark individual T cells that entered the field of view over time.

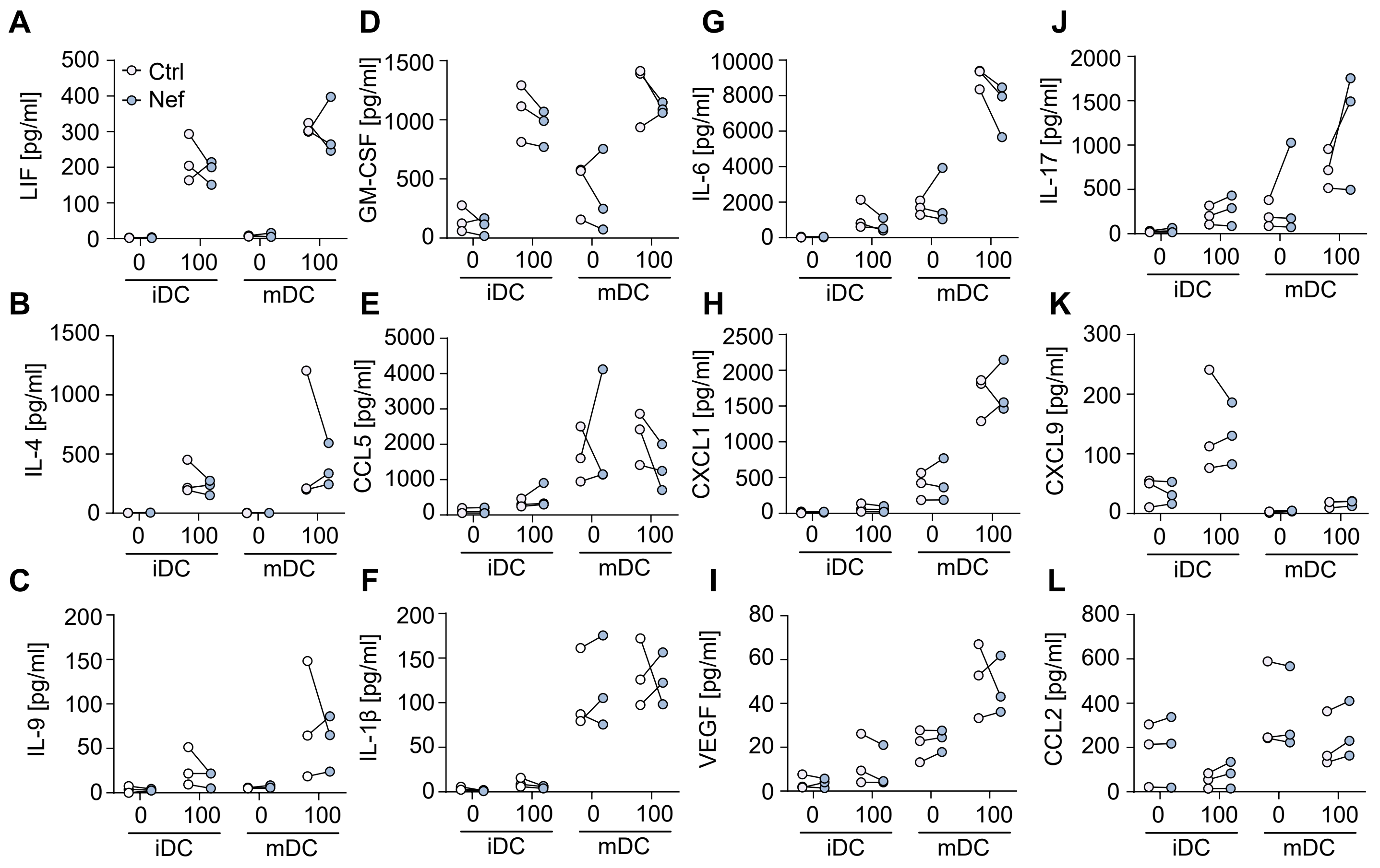


Morath et al. 2025_Fig. S3 for Fig. 3

**Fig. S3. HIV-1 Nef has no effect on a range of cytokines and chemokines related to DC maturation and antigen-specific interaction in T cell co-cultures with DCs. (A)-(L)** Quantification of absolute cytokine concentrations in supernatants of T cell-DC co-cultures, displayed in pg/ml. **(A)-(D)** Cytokines induced in an antigen-specific manner in co-cultures with iDC and mDC (LIF, IL-4, IL-9 , GM-CSF). **(E)-(K)** Cytokines associated with DC maturation status and induced in an antigen-specific manner (CCL5, IL-1β, IL-6, CXCL1, VEGF, IL-17, CXCL9). **(L)** Cytokines without clear association with DC maturation or antigen-specific interaction (CCL2). Shown are values from three independent experiments, each symbol represents data from one experiment and lines connect matched data for Ctrl or Nef co-cultures per experiment. Statistical analysis was performed using Wilcoxon test for LIF, IL-4, IL-9, GM-CSF, CCL5, IL-6, CXCL9 and CCL2 and paired t-test for IL-1β, CXCL1, VEGF and IL-17. p-values are indicated where statistically significant differences between Ctrl and Nef conditions were observed.

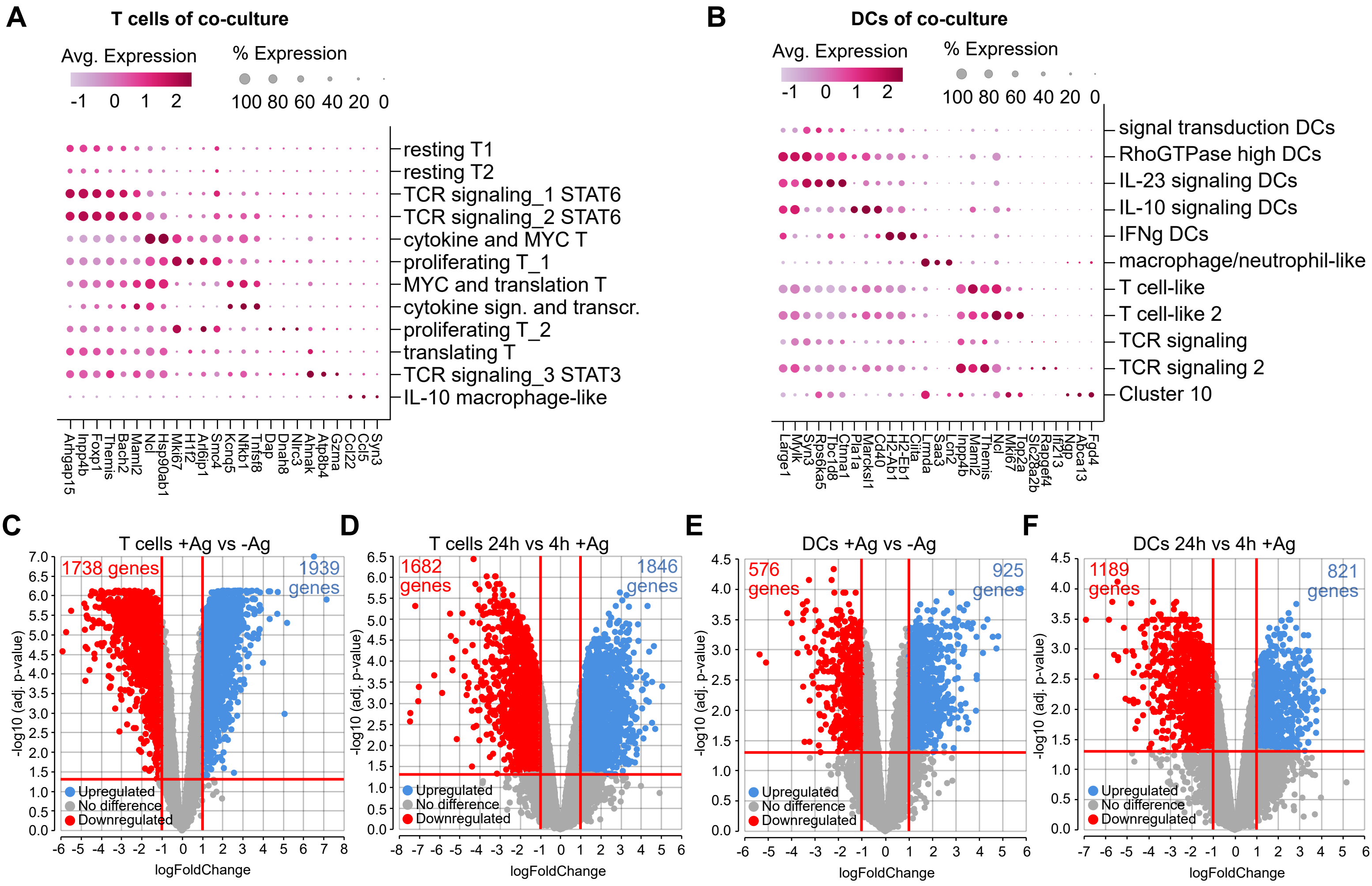
A
T cells of co-culture
Avg. Expression
-1 0 1 2
% Expression
100 80 60 40 20 0
resting T1
resting T2
TCR signaling_1 STAT6
TCR signaling_2 STAT6
cytokine and MYC T
proliferating T_1
MYC and translation T
cytokine sign. and transcr.
proliferating T_2
translating T
TCR signaling_3 STAT3
IL-10 macrophage-like
Arhgap15 Inpp4b Foxp1 Themis Bach2 Maml2 Ncl Hsp90ab1 Mki67 H1f2 Arl6ip1 Smc4 Kcnq5 Nfkb1 Tnfsf8 Dap Dnah8 Nlrc3 Ahnak Atp8b4 Gzma Ccl22 Ccl5 Syn3
B
DCs of co-culture
Avg. Expression
-1 0 1 2
% Expression
100 80 60 40 20 0
signal transduction DCs
RhoGTPase high DCs
IL-23 signaling DCs
IL-10 signaling DCs
IFNg DCs
macrophage/neutrophil-like
T cell-like
T cell-like 2
TCR signaling
TCR signaling 2
Cluster 10
Large1 Mylk Syn3 Rps6ka5 Tbc1d8 Ctnna1 Pla1a Marcksl1 Cd40 H2-Ab1 H2-Eb1 Ciita Lrmda Saa3 Lcn2 Inpp4b Maml2 Themis Ncl Mki67 Top2a Slc28a2b Rapgef4 Ifi213 Ngp Abca13 Fgd4
C
T cells +Ag vs -Ag
1738 genes
1939 genes
Upregulated
No difference
Downregulated
-log10 (adj. p-value)
logFoldChange
D
T cells 24h vs 4h +Ag
1682 genes
1846 genes
Upregulated
No difference
Downregulated
-log10 (adj. p-value)
logFoldChange
E
DCs +Ag vs -Ag
576 genes
925 genes
Upregulated
No difference
Downregulated
-log10 (adj. p-value)
logFoldChange
F
DCs 24h vs 4h +Ag
1189 genes
821 genes
Upregulated
No difference
Downregulated
-log10 (adj. p-value)
logFoldChange

**Fig. S4. Antigen-specific interaction of T cells and DCs induces broad and dynamic transcriptional changes. (A)** and **(B)** Dot plots showing marker gene expression of subclusters within T cells (A) and DCs (B) subgated from total cells in co-culture. **(C)-(F)** Volcano plots showing differential gene expression induced upon antigen-specific interaction in T cell (C, D) or DC (E, F) populations at 4 and 24h timepoints. Shown are comparisons of conditions with or without antigen across both 4 and 24h timepoints for T cells (C) and DCs (E) or comparison of 4 and 24h timepoints of conditions with antigen, for T cells (D) and DCs (F), respectively. Statistical analysis was performed in Trailmaker™ software from Parse Biosciences applying an inbuilt pseudobulk limma-voom workflow for comparisons between groups.

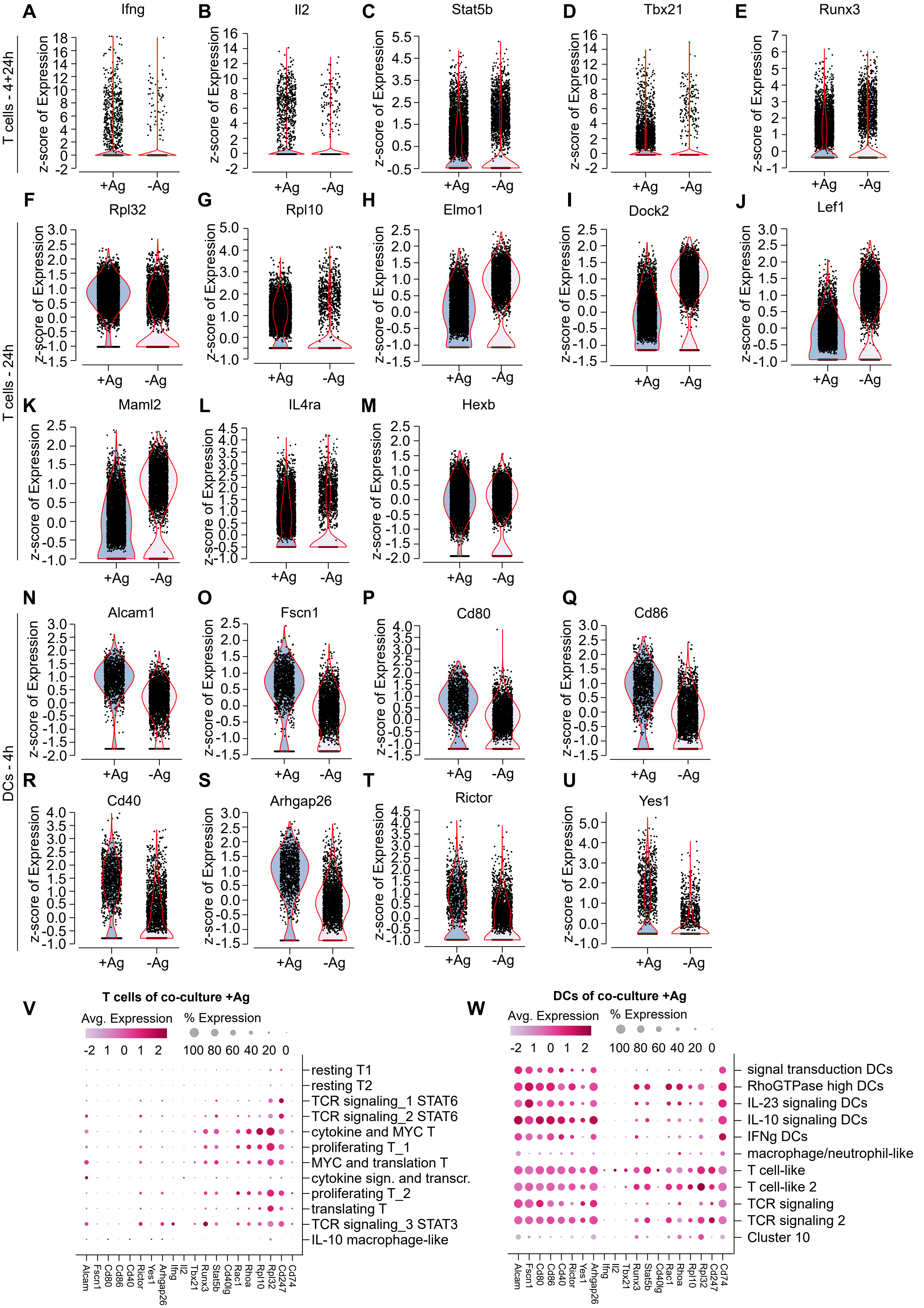


Morath et al. 2025_Fig. S5 for Fig. 4

**Fig. S5. Antigen-specific interaction between T cells and DCs induces expression of activation and maturation-associated genes in T cells and DCs. (A)-(U)** Differentially expressed genes in T cells (A-M) or DCs (N-U) induced by antigen-specific interaction with DCs or T cells, respectively. Shown are violin plots with values for individual cells in samples with (+Ag) or without (-Ag) antigen across both 4 and 24h (4+24h) or at specific timepoints as indicated. **(A)-(E)** Genes upregulated in T cells at both 4 and 24h by antigen-specific interaction with DCs. **(F)** and **(G)** Genes upregulated in T cells at 24h of antigen-specific interaction. **(H)-(M)** Genes repressed (H-K) or induced (L and M) in T cells at 24h of antigen-specific interaction with DCs. **(N)-(U)** Genes upregulated in DCs at 4h of antigen-specific interaction with T cells. A-U. Data is pooled from both Ctrl and Nef samples in co-culture with iDC (24h) or mDC (4+24h). A-E. Data is derived from 6 samples per condition (Ctrl and Nef+iDC 24h and Ctrl and Nef+mDC 4 and 24h). F-M. Data is derived from 4 samples per condition (Ctrl and Nef+iDC 24h and Ctrl and Nef+mDC 24h). N-U. Data is derived from 2 samples per condition (Ctrl and Nef +mDC 4h). **(V)** and **(W)** Dot plots showing magnitude and distribution of expression of antigen-induced genes in the different subclusters identified within UMAPs for T cells (V) or DCs (W).

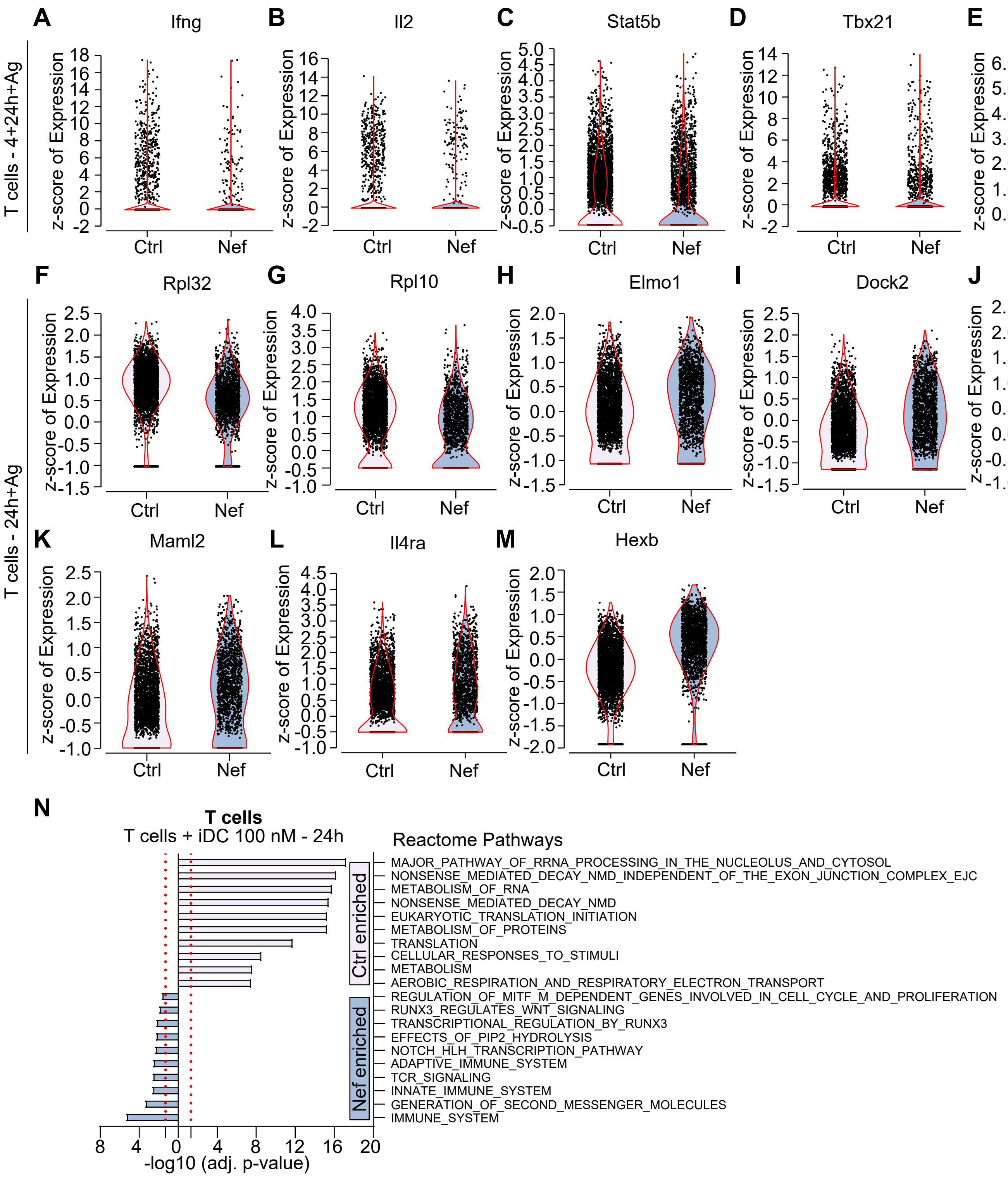
A
Ifng
B
Il2
C
Stat5b
D
Tbx21
E
Runx3
T cells - 4+24h+Ag
z-score of Expression
Ctrl
Nef
F
Rpl32
G
Rpl10
H
Elmo1
I
Dock2
J
Lef1
K
Maml2
L
Il4ra
M
Hexb
T cells - 24h+Ag
N
T cells
T cells + iDC 100 nM - 24h
Reactome Pathways
Ctrl enriched
Nef enriched
MAJOR_PATHWAY_OF_RRNA_PROCESSING_IN_THE_NUCLEOLUS_AND_CYTOSOL
NONSENSE_MEDIATED_DECAY_NMD_INDEPENDENT_OF_THE_EXON_JUNCTION_COMPLEX_EJC
METABOLISM_OF_RNA
NONSENSE_MEDIATED_DECAY_NMD
EUKARYOTIC_TRANSLATION_INITIATION
METABOLISM_OF_PROTEINS
TRANSLATION
CELLULAR_RESPONSES_TO_STIMULI
METABOLISM
AEROBIC_RESPIRATION_AND_RESPIRATORY_ELECTRON_TRANSPORT
REGULATION_OF_MITF_M_DEPENDENT_GENES_INVOLVED_IN_CELL_CYCLE_AND_PROLIFERATION
RUNX3_REGULATES_WNT_SIGNALING
TRANSCRIPTIONAL_REGULATION_BY_RUNX3
EFFECTS_OF_PIP2_HYDROLYSIS
NOTCH_HLH_TRANSCRIPTION_PATHWAY
ADAPTIVE_IMMUNE_SYSTEM
TCR_SIGNALING
INNATE_IMMUNE_SYSTEM
GENERATION_OF_SECOND_MESSENGER_MOLECULES
IMMUNE_SYSTEM
-log10 (adj. p-value)

**Fig. S6. HIV-1 Nef manipulates the transcriptional response in T cells upon antigen-specific interaction with DCs. (A)-(M)** Violin plots showing differentially expressed genes repressed or enhanced by Nef in T cells upon antigen-specific interaction with DCs. **(A)-(E)** Genes repressed by Nef at 4+24h timepoints. **(F)** and **(G)** Genes repressed by Nef at 24h timepoint. **(H)-(M)** Genes enhanced by Nef at the 24h timepoint. Data in A-E is derived from 3 samples per condition (Ctrl or Nef+iDC 24h and Ctrl or Nef+mDC 4 and 24h), data in F-M is derived from 2 samples per condition (Ctrl or Nef+iDC 24h and Ctrl or Nef+mDC 24h). **(N)** Pathway enrichment analysis for subgated T cells comparing Ctrl versus Nef co-cultures with iDC with antigen at 24h using Mouse Reactome 2024 database in MSigDb for enrichment analysis with the top 100 differentially expressed genes. Shown are –log10 values of adjusted p-values. Red dashed lines represent adjusted p-value cutoff of 0.05.

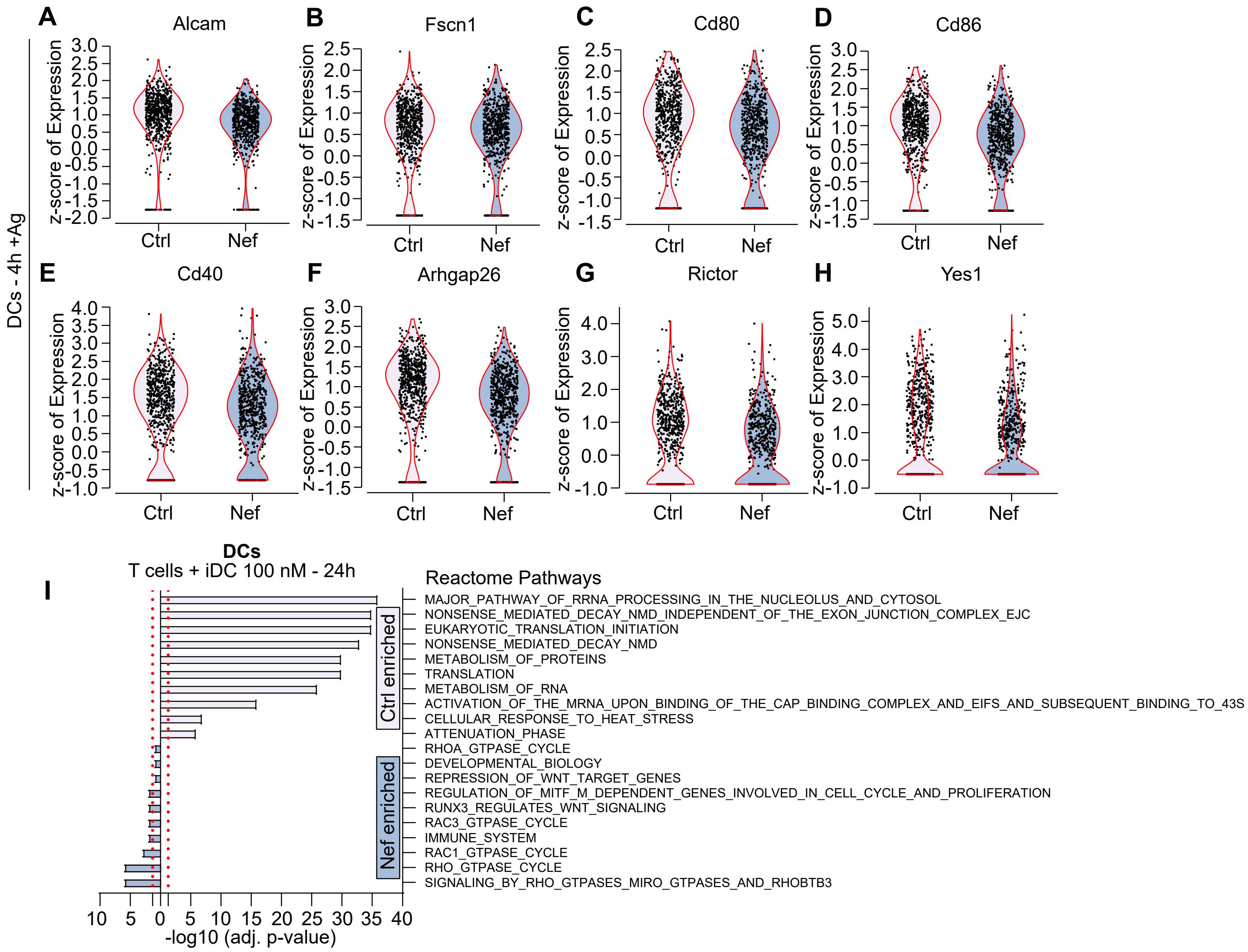
A
Alcam
B
Fscn1
C
Cd80
D
Cd86
E
Cd40
F
Arhgap26
G
Rictor
H
Yes1
DCs - 4h +Ag
z-score of Expression
Ctrl
Nef
I
DCs
T cells + iDC 100 nM - 24h
Reactome Pathways
Ctrl enriched
Nef enriched
MAJOR_PATHWAY_OF_RRNA_PROCESSING_IN_THE_NUCLEOLUS_AND_CYTOSOL
NONSENSE_MEDIATED_DECAY_NMD_INDEPENDENT_OF_THE_EXON_JUNCTION_COMPLEX_EJC
EUKARYOTIC_TRANSLATION_INITIATION
NONSENSE_MEDIATED_DECAY_NMD
METABOLISM_OF_PROTEINS
TRANSLATION
METABOLISM_OF_RNA
ACTIVATION_OF_THE_MRNA_UPON_BINDING_OF_THE_CAP_BINDING_COMPLEX_AND_EIFS_AND_SUBSEQUENT_BINDING_TO_43S
CELLULAR_RESPONSE_TO_HEAT_STRESS
ATTENUATION_PHASE
RHOA_GTPASE_CYCLE
DEVELOPMENTAL_BIOLOGY
REPRESSION_OF_WNT_TARGET_GENES
REGULATION_OF_MITF_M_DEPENDENT_GENES_INVOLVED_IN_CELL_CYCLE_AND_PROLIFERATION
RUNX3_REGULATES_WNT_SIGNALING
RAC3_GTPASE_CYCLE
IMMUNE_SYSTEM
RAC1_GTPASE_CYCLE
RHO_GTPASE_CYCLE
SIGNALING_BY_RHO_GTPASES_MIRO_GTPASES_AND_RHOBTB3
10 5 0 5 10 15 20 25 30 35 40
-log10 (adj. p-value)

**Fig. S7. HIV-1 Nef impairs the transcriptional response in DCs upon antigen-specific interaction with T cells**. **(A)-(H)** Violin plots showing differential gene expression of genes repressed in mDCs upon antigen-specific interaction with Nef T cells compared to Ctrl cells at 4h. Data is derived from one sample per condition (Ctrl or Nef+mDC 4h). **(I)** Pathway enrichment analysis for subgated DCs comparing Ctrl versus Nef co-cultures with iDC with antigen at 24h using Mouse Reactome 2024 database in MSigDb for enrichment analysis with the top 100 differentially expressed genes. Shown are –log10 values of adjusted p-values. Red dashed lines represent adjusted p-value cutoff of 0.05.

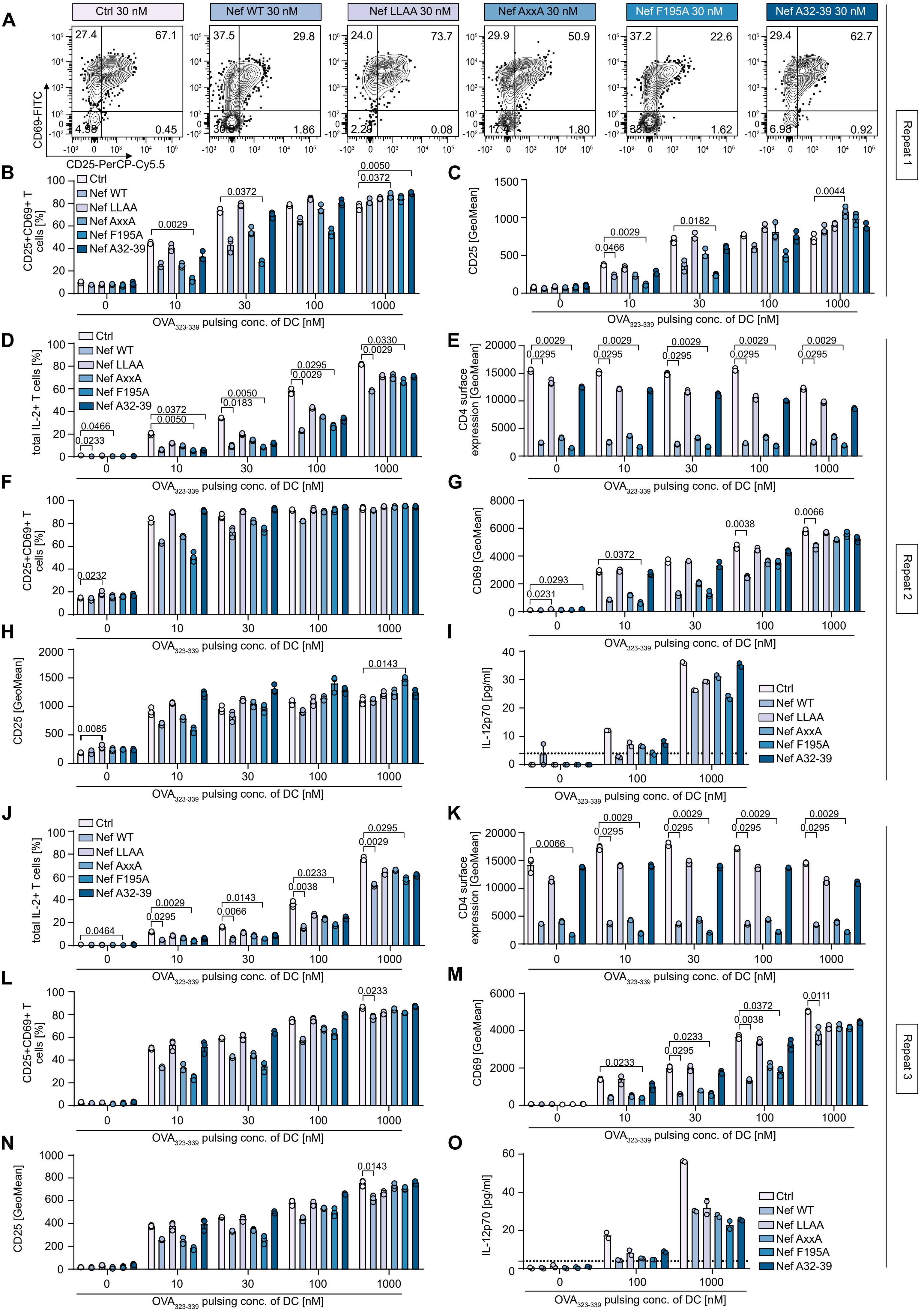


Morath et al. 2025_Fig. S8 for Fig. 5

**Fig. S8. The di-leucine motif and S5AM of HIV-1 Nef reproducibly confer interference with CD4 T cell-DC interactions. (A)-(C)** Supplementary data obtained from the experiment in Fig. 5. **(A)** Representative flow cytometry contour plots for the indicated conditions showing CD25 and CD69 expression upon co-culture with mDCs pulsed with 30 nM OVA antigen at 24h. **(B)** Quantification of A. **(C)** Quantification of CD25 GeoMean on CD4 T cells for conditions in A and B. **(D)-(O)** Additional independent experimental repeats to Fig. 5 showing IL-2 production (D and J), CD4 surface expression (E and K) and CD25/CD69 upregulation (F-H and L-N) of OT-II.2 CD4 T cells transduced with Ctrl or SF2 Nef or mutant MLV in co-cultures with mDCs at indicated peptide pulsing concentrations. **(I)** and **(O)** IL-12p70 concentrations measured in supernatants of 24h co-cultures from F-H or L-N by ELISA, respectively. Dashed line indicates the detection limit for the assay. Shown are means with SD of biological triplicates, i.e. technical duplicates with range for I and O. Statistical analysis was performed using Kruskal-Wallis test, per condition, SF2 Nef and mutant CD4 T cell conditions were compared with the Ctrl CD4 T cell condition only. p-values are indicated for statistically significant differences.

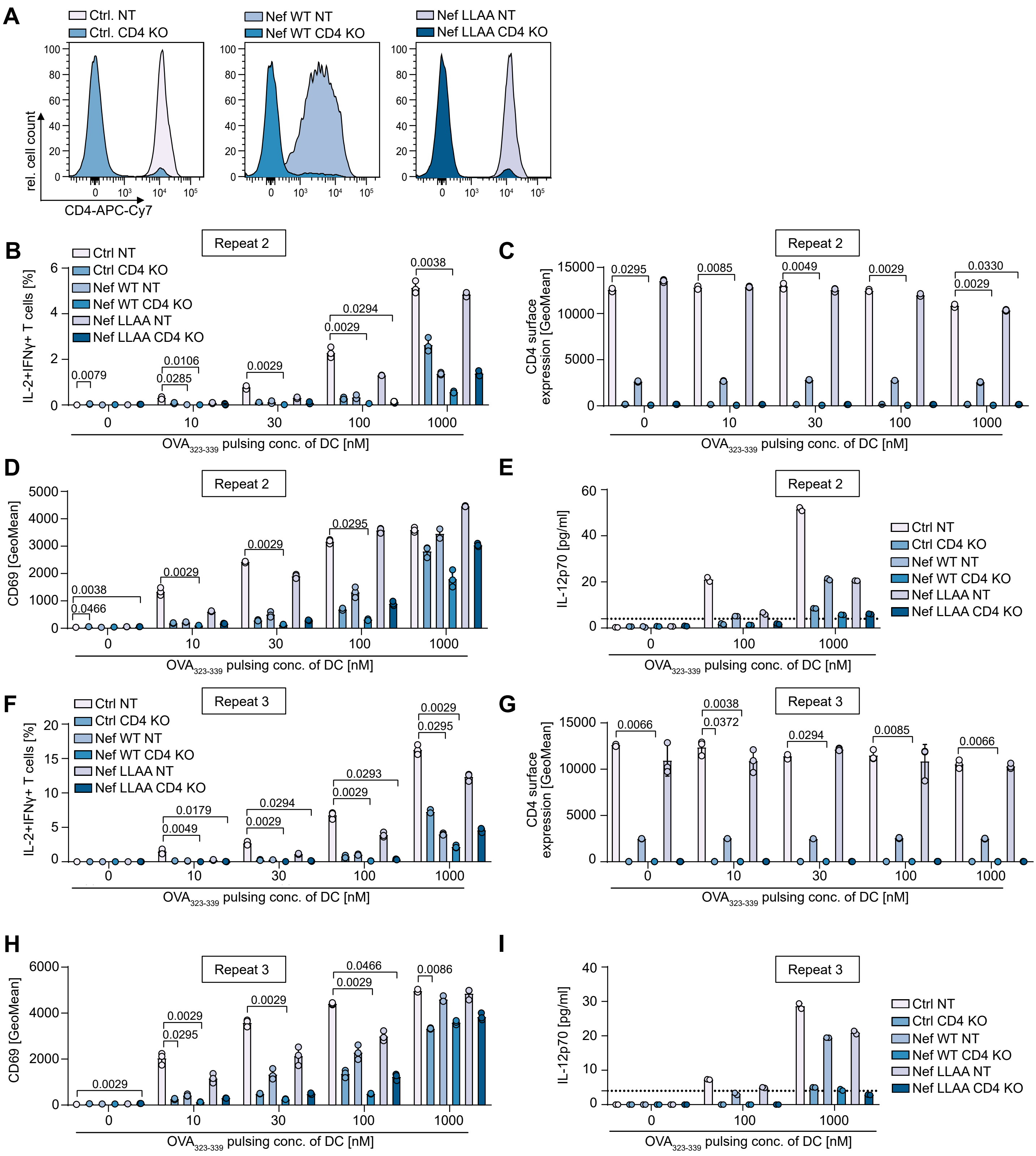
A
Ctrl. NT
Ctrl. CD4 KO
Nef WT NT
Nef WT CD4 KO
Nef LLAA NT
Nef LLAA CD4 KO
rel. cell count
CD4-APC-Cy7
B
Repeat 2
IL-2+IFNγ+ T cells [%]
OVA323-339 pulsing conc. of DC [nM]
C
CD4 surface expression [GeoMean]
D
CD69 [GeoMean]
E
IL-12p70 [pg/ml]
F
Repeat 3
G
H
I
Ctrl NT
Ctrl CD4 KO
Nef WT NT
Nef WT CD4 KO
Nef LLAA NT
Nef LLAA CD4 KO

**Fig. S9. HIV-1 Nef's interference with CD4 T cell interaction with DCs reproducibly depends on its CD4 downmodulation function. (A)** Representative flow cytometry histograms showing CD4 KO efficiency in NGFR-sorted OT-II.2 cells in co-culture with mDC with 1000 nM antigen on day 8. Shown are superimposed signals for Ctrl (left), Nef WT (middle) or Nef LLAA (right) cells nucleofected with NT or CD4 RNP as indicated. **(B)-(I)** Additional independent experimental repeats to Fig. 6. **(B)** and **(F)** Quantification of cytokine production in OT-II.2 CD4 T cells nucleofected with NT or CD4 KO RNP upon cognate co-culture with mDCs at 4.5h. **(C)** and **(G)** Quantification of surface CD4 levels on OT-II.2 cells for conditions from B and F. **(D)** and **(H)** Quantification of surface CD69 levels on CD4 T cells from indicated conditions at 24h of co-culture with peptide-pulsed mDCs. **(E)** and **(I)** Quantification of IL-12p70 levels in supernatants of indicated conditions of CD4 T cells in mDC co-cultures at 24h measured by ELISA. Dashed line indicates the detection limit for the assay. Shown are means with SD of biological triplicates, i.e. technical duplicates for E and I. Statistical analysis was performed using Kruskal-Wallis test, per peptide-pulsing condition, SF2 Nef or Nef LLAA mutant with NT or CD4 KO RNP nucleofection were compared with the Ctrl CD4 T cell NT condition. p-values are indicated for statistically significant differences.

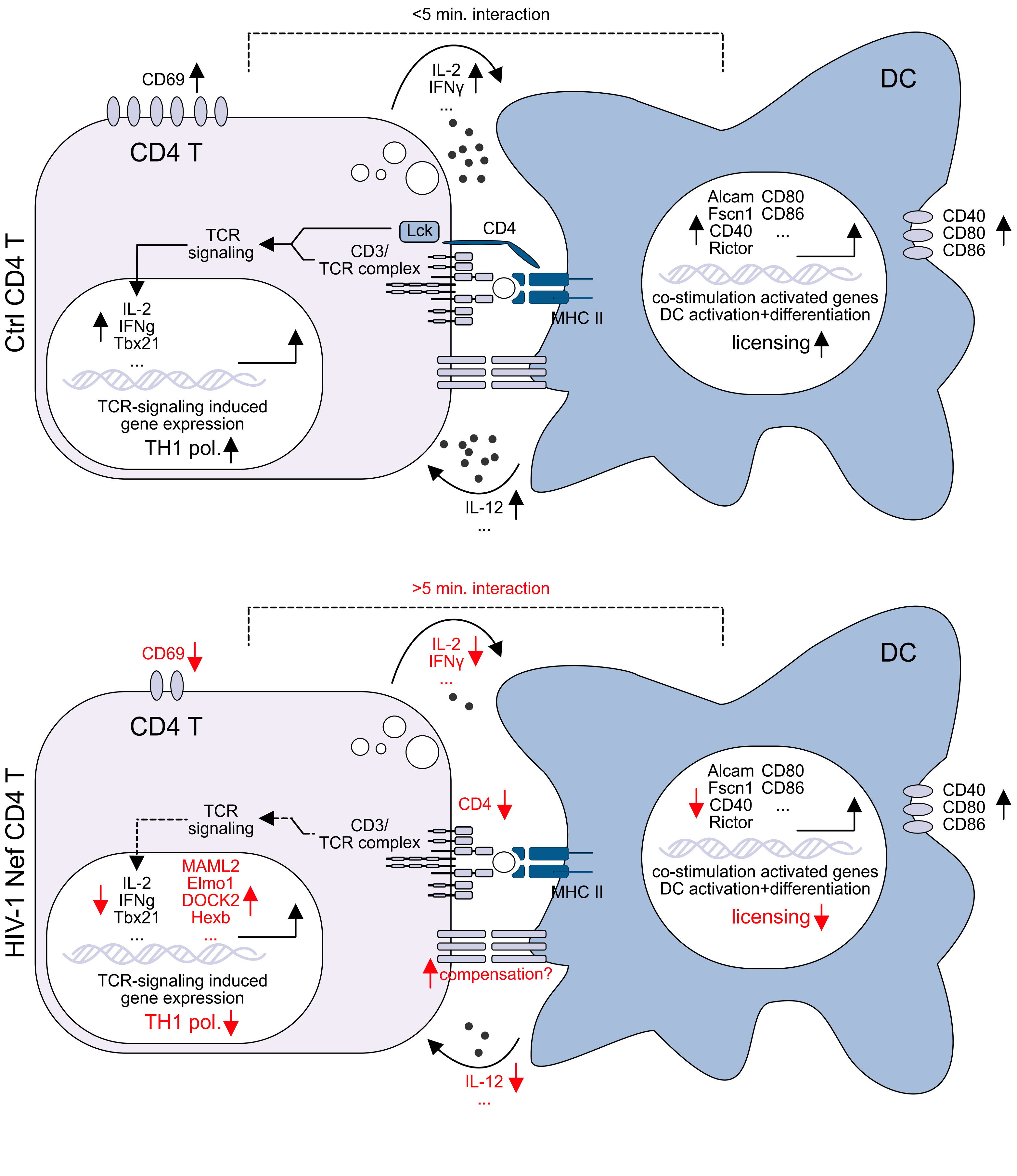


Morath et al. 2025_Fig. S10

**Fig. S10. Graphical abstract visualizing manipulation of cognate CD4 T cell-DC interaction by HIV-1 Nef.** By downmodulation of CD4, Nef manipulates T cell-DC interactions, leading to inefficient antigen-specific activation of both T cells and DCs that is associated with a transcriptional reprograming of CD4 T cells, disfavoring Th1 polarization.

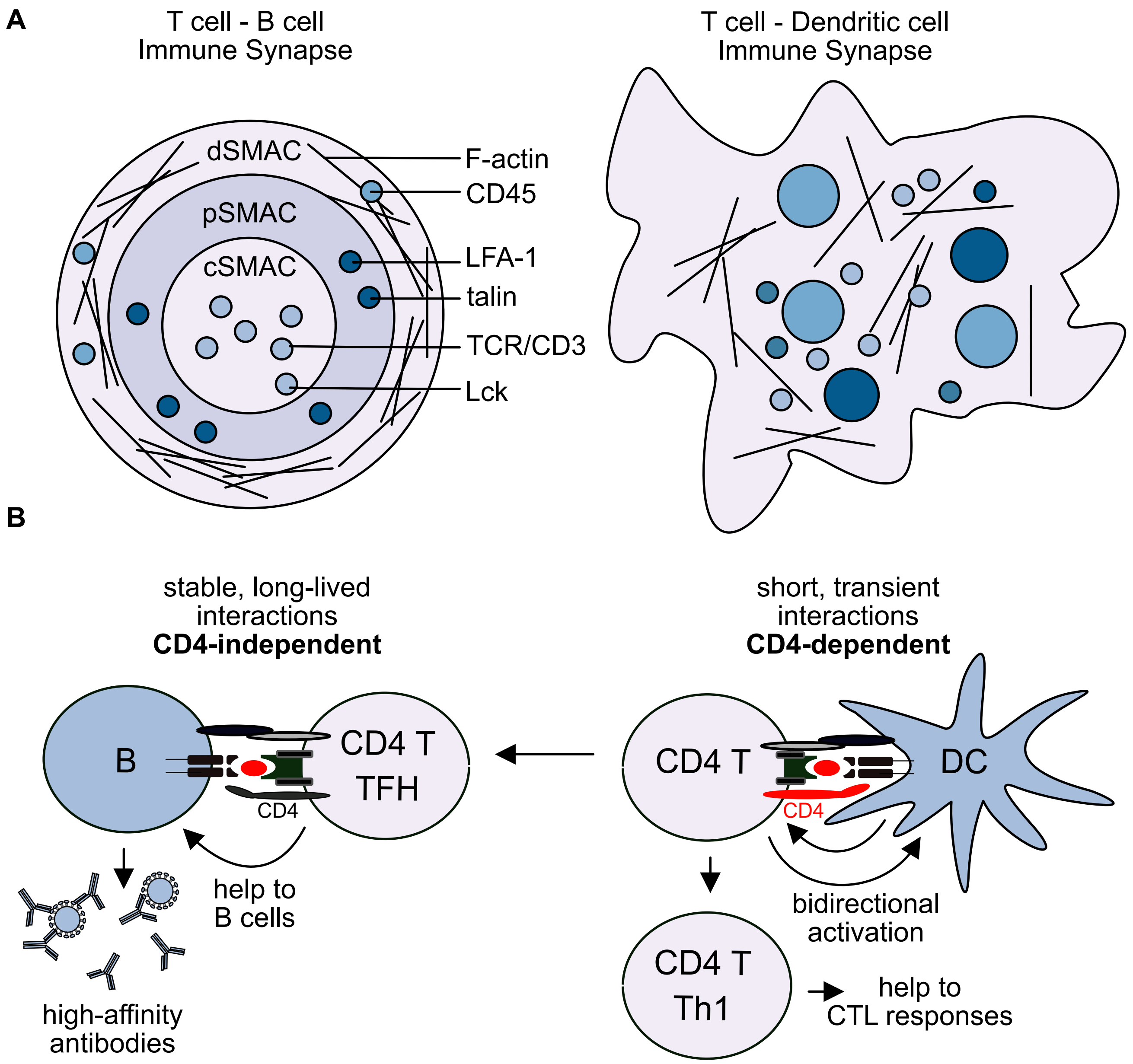


Morath et al. 2025_Fig. S11

**Fig. S11. Schematic comparison of the architecture and dynamics of ISes of CD4 T cells with B cells or with DCs, respectively**. **(A)** While CD4 T cell synapses with B cells are highly organized into supramolecular activation clusters (SMACs), CD4 T cell synapses with DCs are multifocal and disorganized. **(B)** T cell-B cell synapses are stable, long-lived and CD4 independent while T cell-DC synapses are highly dynamic, transient and CD4 dependent.

| Sample ID | Experimental repeat | Condition |
|---|---|---|
| S1 | 1 | Ctrl + iDC 0 nM OVA |
| S2 | 1 | Ctrl + iDC 100 nM OVA |
| S3 | 1 | Ctrl + mDC 0 nM OVA |
| S4 | 1 | Ctrl + mDC 100 nM OVA |
| S5 | 1 | Nef + iDC 0 nM OVA |
| S6 | 1 | Nef + iDC 100 nM OVA |
| S7 | 1 | Nef + mDC 0 nM OVA |
| S8 | 1 | Nef + mDC 100 nM OVA |
| S9 | 2 | Ctrl + iDC 0 nM OVA |
| S10 | 2 | Ctrl + iDC 100 nM OVA |
| S11 | 2 | Ctrl + mDC 0 nM OVA |
| S12 | 2 | Ctrl + mDC 100 nM OVA |
| S13 | 2 | Nef + iDC 0 nM OVA |
| S14 | 2 | Nef + iDC 100 nM OVA |
| S15 | 2 | Nef + mDC 0 nM OVA |
| S16 | 2 | Nef + mDC 100 nM OVA |
| S17 | 3 | Ctrl + iDC 0 nM OVA |
| S18 | 3 | Ctrl + iDC 100 nM OVA |
| S19 | 3 | Ctrl + mDC 0 nM OVA |
| S20 | 3 | Ctrl + mDC 100 nM OVA |
| S21 | 3 | Nef + iDC 0 nM OVA |
| S22 | 3 | Nef + iDC 100 nM OVA |
| S23 | 3 | Nef + mDC 0 nM OVA |

| | | |
|---|---|---|
| S24 | 3 | Nef + mDC 100 nM OVA |

**Table S1 for Data S1: Sample key for Eve technologies Mouse Cytokine 32-Plex Discovery Assay**

Sample key for analysis of cytokines in ex vivo co-cultures of OT-II.2 CD4 T and iDC or mDC pulsed with or without 100 nM OVA323-339. Only cytokines in which values for all three experimental repeats could be determined were included for final analysis.

| Commercial reagents | | |
|---|---|---|
| **Item** | **Manufacturer** | **Cat. Nr.** |
| DMEM high Glucose | Thermo Fisher Scientific | 61965-026 |
| RPMI 1640 | Thermo Fisher Scientific | 61870-010 |
| Penicillin/Streptomycin | Thermo Fisher Scientific | 15140-122 |
| FBS | Capricorn Scientific | FBS-HI-11A |
| Non-essential amino acids | Thermo Fisher Scientific | 11140-035 |
| Sodium pyruvate | Thermo Fisher Scientific | 11360-039 |
| β-Mercaptoethanol | Thermo Fisher Scientific | 21985-023 |
| RetroNectin® Recombinant Human Fibronectin Fragment | Takara Bio | T100A |
| OVA323-339 | InvivoGen | vac-isq |
| Mouse IL-7 Recombinant Protein, PeproTech® | Thermo Fisher Scientific | 217-17-10UG |
| Recombinant Mouse GM-CSF (carrier-free) | BioLegend | 576304 |
| Puromycin | Calbiochem | 540411 |
| Blasticidin S HCl, Pulver | Thermo Fisher Scientific | R21001 |
| Monensin Solution (1,000X) | BioLegend | 420701 |
| JetPEI™ | Polyplus | 101000020 |
| OptiPrep™ | Stemcell technologies | 07820 |
| Collagenase/Dispase® | Merck | 10269638001 |
| Alt-R™ S.p. Cas9 Nuclease v3 | Integrated DNA technologies (IDT) | 1081059 |

| | | |
|---|---|---|
| FTY720 | Sigma Aldrich | SML0700 |
| Lipopolysaccharide from E. coli | Sigma Aldrich | L2654-1MG |
| **Commercial kits** | | |
| **Item** | **Manufacturer** | **Cat. Nr.** |
| EasySep™ Human CD271 Positive Selection Kit II | Stemcell technologies | 17849 |
| Evercode™ WT chemistry version 3 kit | Parse Biosciences | UMWT3300 |
| Evercode™ Cell Fixation v3 kit | Parse Biosciences | ECFC3300 |
| P3 Primary Cell 4D-nucleofector™ X kit, L | Lonza | V4XP-3024 |
| ELISA MAX™ Deluxe Set Mouse IL-12 (p70) | BioLegend | 433604 |
| BD Cytofix/Cytoperm™ Fixation/Permeabilization Kit | BD Biosciences | 554714 |
| True-Nuclear™ Transcription Factor Buffer Set | BioLegend | 424401 |
| Zombie Violet™ Fixable Viability Kit | BioLegend | 423113 |
| CellTrace™ CFSE Cell Proliferation Kit | Thermo Fisher Scientific | C34570 |

| | | |
|---|---|---|
| CellTrace™ Violet Cell Proliferation Kit | Thermo Fisher Scientific | C34557 |
| AlexaFluor 647 NHS-Ester | Thermo Fisher Scientific | A20006 |
| **House-made solutions** | | |
| **Item** | **Components** | |
| CMR | RPMI 1640 + 10% FBS + 1% Penicillin/Streptomycin + 10 mM HEPES + 1% non-essential amino acids + 1% sodium pyruvate + 50 µM β-Mercaptoethanol | |
| ACK buffer | Pre-mixed 8,29g $NH_4Cl$ + 1g $KHCO_3$ in 500ml $H_2O$ millipore combined with pre-mixed 0,0367g EDTA + 100ml $H_2O$ millipore, filled up to 1l, pH adjusted (pH=7.6) and sterile filtered | |
| BMDC medium | RPMI 1640 + 10% FBS +1% Penicillin/Streptomycin + 50 µM β-Mercaptoethanol | |
| FACS buffer | 1xPBS + 0.5% BSA +2 mM EDTA | |
| Recomm. Medium for NGFR sorting | 1xPBS + 2mM EDTA + 2% FBS | |
| **Relevant consumables** | | |
| **Item** | **Manufacturer** | **Cat. Nr.** |
| 0.45 µm filter stericup | Merck | S2HVU05RE |
| 0.45 µm syringe filter | Carl ROTH | KH55.1 |
| 27G Microlance needle | BD Biosciences | 302200 |
| pluriStrainer 70µm, PET-Mesh | Pluriselect | 43-50070-51 |

| | | |
|---|---|---|
| pluristrainer 100 µm, PET-Mesh | Pluriselect | 43-50100-51 |
| Petri dishes for BMDCs | Greiner | 664102 |
| 96-well NUNC™ Maxisorp plates (ELISA) | Thermo Fisher Scientific | 442404 |

**sgRNAs**

| **Target** | **Sequence (5’-3’)** | **Manufactuer** |
|---|---|---|
| Murine CD4_1 | AACUCCUAGCUGUCACUCAA | Synthego |
| Murine CD4_2 | UCAAAACGAUCAAACUGCGA | Synthego |
| Murine CD4_3 | UUCUUCUGGGAACUCUCGCA | Synthego |

**Antibodies**

| **Item** | **Manufacturer** | **Clone** | **Cat. Nr.** | **RRID** |
|---|---|---|---|---|
| CD3-APC-Cy7 | BioLegend | 17A2 | 100222 | AB_2242784 |
| CD45.2-PE | BioLegend | 104 | 109808 | AB_313445 |
| CD69-FITC | BioLegend | H1.2F3 | 104506 | AB_313109 |
| NGFR-PE | BioLegend | ME20.4 | 345106 | AB_2152647 |
| CD11c-APC | BioLegend | N418 | 117310 | AB_313779 |
| CD25-PerCP-Cy5.5 | BioLegend | 3C7 | 101912 | AB_10613643 |
| CD69-BV650 | BioLegend | H1.2F3 | 104541 | AB_2616934 |
| CD86-PerCP | BioLegend | GL-1 | 105026 | AB_893417 |
| CD4-APC-Cy7 | BioLegend | GK1.5 | 100414 | AB_312699 |
| IFNγ-FITC | BioLegend | XMG1.2 | 505806 | AB_315400 |
| IL-2-APC | BioLegend | JES6-5H4 | 503810 | AB_315304 |

| | | | | |
|---|---|---|---|---|
| T-bet-PerCP-Cy5.5 | BioLegend | 4B10 | 644806 | AB_1595488 |
| TruStain FcX™ (anti-mouse CD16/32) Antibody | BioLegend | 93 | 101319 | AB_1574973 |
| Meca-79-AF647 | NanoTools | Meca-79 | Custom made | |
| Meca79-AF647 | Santa Cruz Biotechnology | Meca-79 | sc-19602 AF647 | AB_627143 |
| **Plasmids** | | | | |
| **Item** | | **Source** | | |
| MLV-A | | Stolp et al., PNAS 2012 | | |
| pSTITCH ΔNGFR | | Stolp et al., PNAS 2012 | | |
| pSTITCH HIV-1 SF2 Nef ΔNGFR | | Stolp et al., PNAS 2012 | | |
| pSTITCH HIV-1 SF2 Nef LLAA | | Kaw et al., EMBO J 2020 | | |
| **Cell lines** | | | | |
| **Item** | | **Source** | | |
| Plat-E MLV packaging cell line | | Morita et al., Gene Therapy 2000 | | |

**Table S2. Supplementary materials list.**

**Data S1. Excel sheet with raw data of Eve technologies Mouse Cytokine 32-Plex Discovery Assay from Eve technologies.**

**Movie S1. Control T cells interacting with 1000 nM OVA peptide-pulsed mBMDCs after 4-12h of interactions**. Shown are OT-II.2cfp Ctrl CD4 T cells in blue, membrane-associated tdTomato (mT) expressing mBMDCs pulsed with 1000 nM OVA peptide in red and high endothelial venules (HEVs), labeled with Meca79-AlexaFluor 647 in grey. Movie was acquired 4-12h post DC arrival in the draining lymph node, major grid and scale bar 40 µm, 121 timesteps of 20 seconds. In the second half of the movie the zoomed in area shown in the still images in Fig. S2C are presented.

**Movie S2. Control T cells interacting with 1000 nM OVA peptide-pulsed mBMDCs after 20-28h of interactions**. Shown are OT-II.2cfp Ctrl CD4 T cells in blue, membrane-associated tdTomato (mT) expressing mBMDCs pulsed with 1000 nM OVA peptide in red and high endothelial venules (HEVs), labeled with Meca79-AlexaFluor 647 in grey. Movie was acquired 20-28h post DC arrival in the draining lymph node, major grid and scale bar 40 µm, 121 timesteps of 20 seconds. In the second half of the movie the zoomed in area shown in the still images in Fig. 2I are presented.

**Movie S3. Migration of Control T cells and unpulsed mBMDCs**. Shown are OT-II.2cfp Ctrl CD4 T cells in blue, membrane-associated tdTomato (mT) expressing mBMDCs in red and high endothelial venules (HEVs), labeled with Meca79-AlexaFluor 647 in grey. Movie was acquired post DC arrival in the draining lymph node, major grid and scale bar 40 µm, 121 timesteps of 20 seconds. In the second half of the movie the zoomed in area shown in the still images in Fig. S2C are presented.

**Movie S4. Nef T cells interacting with 1000 nM OVA peptide-pulsed mBMDCs after 4-12h of interactions**. Shown are OT-II.2cfp Nef CD4 T cells in blue, membrane-associated tdTomato (mT) expressing mBMDCs pulsed with 1000 nM OVA peptide in red and high endothelial venules (HEVs), labeled with Meca79-AlexaFluor 647 in grey. Movie was acquired 4-12h post DC arrival in the draining lymph node, major grid and scale bar 40 µm, 121 timesteps of 20 seconds. In the second half of the movie the zoomed in area shown in the still images in Fig. S2C are presented.

**Movie S5. Nef T cells interacting with 1000 nM OVA peptide-pulsed mBMDCs after 20-28h of interactions**. Shown are OT-II.2cfp Nef CD4 T cells in blue, membrane-associated tdTomato (mT) expressing mBMDCs pulsed with 1000 nM OVA peptide in red and high endothelial venules (HEVs), labeled with Meca79-AlexaFluor 647 in grey. Movie was acquired 20-28h post DC arrival in the draining lymph node, major grid and scale bar 40 µm, 121 timesteps of 20 seconds. In the second half of the movie the zoomed in area shown in the still images in Fig. 2I are presented.

**Movie S6. Migration of Nef T cells and unpulsed mBMDCs**. Shown are OT-II.2cfp Nef CD4 T cells in blue, membrane-associated tdTomato (mT) expressing mBMDCs in red and high endothelial venules (HEVs), labeled with Meca79-AlexaFluor 647 in grey. Movie was acquired post DC arrival in the draining lymph node, major grid and scale bar 40 µm, 121 timesteps of 20 seconds. In the second half of the movie the zoomed in area shown in the still images in Fig. S2C are presented.
